# Discovery of a pre-mRNA structural scaffold as a contributor to the mammalian splicing code

**DOI:** 10.1101/292458

**Authors:** Kaushik Saha, Mike Minh Fernandez, Tapan Biswas, Simpson Joseph, Gourisankar Ghosh

## Abstract

The specific recognition of splice signals at or near exon-intron junctions is not explained by their weak conservation and instead is postulated to require a multitude of features embedded in the pre-mRNA strand. We explored the possibility of three-dimensional structural scaffold of *AdML* – a model pre-mRNA substrate – guiding early spliceosomal components to the splice signal sequences. We find that mutations in the non-cognate splice signal sequences impede recruitment of early spliceosomal components due to disruption of the global structure of the pre-mRNA. We further find that the pre-mRNA segments potentially interacting with the early spliceosomal component U1 snRNP are distributed across the intron, that there is a spatial proximity of 5′ and 3′ splice sites within the pre-mRNA scaffold, and that an interplay exists between the structural scaffold and splicing regulatory elements in recruiting early spliceosomal components. These results suggest that early spliceosomal components can recognize a three-dimensional structural scaffold beyond the short splice signal sequences, and that in our model pre-mRNA, this scaffold is formed across the intron involving the major splice signals. This provides a conceptual basis to analyze the contribution of recognizable three-dimensional structural scaffolds to the splicing code across the mammalian transcriptome.

## INTRODUCTION

The early spliceosome defines the exon-intron boundaries within pre-mRNAs for eventual removal of introns and ligation of exons by the mature spliceosome. The early spliceosome assembles at the exon-intron junction via stepwise recognition of multiple splice signals on pre-mRNAs by a set of highly conserved core spliceosomal components (1). The four major splice signals, namely 5′ and 3′ splice sites (SS) located at 5′ and 3′ ends of the intron, the branch-point site (BS) located approximately 30 to 50 nucleotides (nt) upstream of the 3′SS, and the polypyrimidine tract (PPT) of varying lengths present between the BS and the 3′SS, are bound by the U1 small nuclear ribonucleoprotein (U1 snRNP), U2 auxiliary factor 35 (U2AF35), splicing factor 1 (SF1), and U2 auxiliary factor 65 (U2AF65), respectively (1). In the early spliceosome, the 5′SS base-pairs with the 5′ end of U1 snRNA, the RNA component of U1 snRNP, generating an RNA-RNA hybrid that can be up to 11-nt in length (2). Splicing can be constitutive, where the same set of splice sites are selected in all molecules of the synthesized transcript, or alternative, where more than one set of splice sites are selected in different cell or tissue types or in the same cell type under different conditions.

Splice signal recognition is a highly efficient process, and errors in splice signal recognition including splicing from unannotated splice sites (cryptic splicing) often lead to diseases (3,4). How the yeast mature spliceosome prevents incorporation of a sub-optimal/cryptic splice site is currently being investigated (5,6). However, mechanisms of efficient and faithful recognition of the enormous combinatorial library of splice signals in the mammalian transcriptome by the early spliceosomal components are still not fully understood. This is primarily because the conserved region of the mammalian splice signals is short, often about two nucleotides; this level of degeneracy within the splice signal sequences often blurs the distinction between the authentic splice signal sequences and random sequences to the common eye. That is why splicing research has been focused on generating an effective ‘splicing code’ that could explain the outcome of a splicing event, particularly where multiple pairs of alternatively selected splice sites are present (7,8). So far, three features of the pre-mRNA are considered to contribute to the ‘splicing code’: the sequence of the splice signals, local and long-distance pre-mRNA secondary structure, and the distribution of splicing regulatory elements (SREs) that could be exonic/intronic enhancers/silencers. SREs recruit from a large repertoire of RNA binding proteins (RBP) for managing the recognition-potential of a splice signal (9). Recently, the use of deep learning technology and mathematical modeling have greatly enhanced our ability to predict splicing outcome using the genomic sequence (4,10). However, many of the mechanistic details remain obscure.

Given that enhancers augment the pre-mRNA features that are recognized by the early spliceosomal components, understanding the mechanism of action of enhancers is important for understanding the splice signal recognition mechanism. The well-studied exonic splicing enhancers (ESE) bind one of numerous RBPs to promote incorporation of a specific splice signal into the early spliceosome, and their functionalities are negatively impacted by the level of base-pairing within the site (11). In order to identify and characterize the potential ESE-regulated splicing events (both constitutive and alternative) in the mammalian transcriptome, the distribution (12–21) and local secondary structural environment (12,18,22) of the ESEs are being intensely investigated. With serine-arginine-rich (SR) proteins (23), which are some of the most well-characterized ESE-recruited RBPs, it has been demonstrated that these ESE-bound RBPs promote splicing by directly interacting with the early spliceosomal component and stabilizing it at the splice signal (24). Recently, it has been reported that assembly of the early spliceosome could also be promoted by structural remodeling of the pre-mRNA mediated by ESE-recruited RBPs (18). Despite significant advancements in our understanding of the ‘splicing code’, the context-dependence of splice signal usage often cannot be mechanistically explained (25). This suggests that the diversity of pre-mRNA features constituting the splicing code is yet to be fully comprehended.

Biochemical and structural studies of the yeast early spliceosomal complex (E-complex) suggest the formation of an early cross-intron bridge between the splice sites (26,27). In the current study, we explored if the global three-dimensional structural scaffold of the protein-free pre-mRNA could mediate cross-intron communication and guide the recruitment of early spliceosomal components. We tested this possibility by examining recruitment of early spliceosomal components to a protein-free splicing substrate *in vitro*. Recruitment of early spliceosomal components strictly dependent on ESEs might complicate the experimental analysis of the effect of the pre-mRNA structure on defining the splice signal. Therefore, we searched for a splicing substrate where U1 snRNP recruitment is sufficiently strong and specific even in the absence of ESE-dependent functionalities, and found the well-studied model pre-mRNA substrate *AdML* (adenovirus 2 major late transcript IVS1) (see Materials and Methods) (28). Mutation in any of the splice signals in *AdML* disrupted its global structure and the recruitment of the early spliceosomal components U1 snRNP, U2AF65, and U2AF35. We also correlated the strandedness and spatial distribution of nucleotides across the protein-free *AdML* pre-mRNA with their potential for interaction with U1 snRNP anchored at the 5′SS. Additionally, we found that the global *AdML* pre-mRNA structure that ‘integrates’ the splice signals also regulates the stabilizing effect of ESE-associated RBPs on early spliceosomal components for binding to the splicing substrate. These results demonstrate the contribution of the global structural landscape of any pre-mRNA tested so far to the mammalian ‘splicing code’, and open up avenues for further investigation into its potential to regulate splicing efficiency.

## MATERIALS AND METHODS

### Cloning and protein expression

The cDNAs of all proteins except for that of full-length U2AF35 were cloned into T7 promoter-based *E. coli* expression vectors and were expressed as either non-fusion or hexa-histidine (His_6_) fusion proteins. Proteins were expressed in *E. coli* BL21 (DE3), BL21 (DE3) pLysS, or Rosetta (DE3) cells overnight without (leaky expression) or with isopropyl β-D-1-thiogalactopyranoside induction. His_6_-tagged proteins were purified by Ni^2+^-nitrilotriacetate (Ni-NTA) chromatography while non-fusion proteins were purified by a combination of SP Sepharose (Cytiva), hydroxyapatite (Bio-Rad), and butyl Sepharose (Cytiva). His_6_-tagged U2AF35 was expressed and purified from baculovirus-infected Sf9 cells as described before (29) and purified by Ni-NTA chromatography under denaturing conditions (8M urea). Then the protein was refolded by removing the urea stepwise from the buffer through multiple rounds of dialysis. All these proteins were further purified by size-exclusion (Superdex 75; Cytiva) chromatography. His_6_-tagged Sm core proteins were co-expressed in combinations (D3-B, D1-D2, and E-F-G) and purified as described before (30). His_6_-U2AF65 was either expressed on its own or co-expressed with His_6_-tagged truncated SF1_320_ (1-320 amino acids) in *E. coli*. After purification, SF1_320_ and U2AF65 could be separated from each other by gel filtration.

Full-length SRSF1 (hyperphosphorylated mimetic with all serine residues of the RS domain [197-246 a.a.] replaced with glutamate (31)) and the RNA binding domain (RBD) of SRSF5 (1-184 a.a.) were used in chromatographic and pull-down assays. The RBD of SRSF1 (1-203 a.a.) was used for EMSA since the full-length SRSF1 precipitates upon contact with the native gel running buffer in the well. All SR proteins were freed from most of the *E. coli* RNA bound to it by mixing the proteins with Q Sepharose. A chimera of maltose binding protein (MBP) and bacteriophage MS2-coat protein (MS2) was expressed in *E. coli* as a His_6_-tagged protein and purified by Ni-NTA chromatography. The chimeric protein migrated as a doublet on SDS gel and protein(s) represented in both bands comprised of fully functional MBP-MS2. GST-tagged proteins used for pull-down assays were purified by Glutathione Sepharose (Cytiva) and then were cleaned up further as indicated above. All purified proteins were confirmed to be RNase-free by incubating a small aliquot of the purified protein with purified U1 snRNA overnight at room temperature and analyzing the RNA quality by urea PAGE following phenol extraction.

### RNA constructs

*AdML* (adenovirus type 2 major late transcript IVS1) sequence was as described before (18). Adenovirus type 2 major late transcript has been used extensively as a model pre-mRNA substrate since the discovery of splicing (32,33). Since then, it has been modified to generate several variants and subjected to numerous mechanistic and structural studies of spliceosome assembly (27,34–38). The variant used in this study was purchased from Addgene and was deposited by Robin Reed (39,40). All pre-mRNAs except for *AdML* EH mutant contained three MS2-coat protein binding loops at their 3′ end as reported before (40). Changes in nucleotide sequences for mutagenesis of splice signals of *AdML*: Δ5′SS (41) (GTTGGGgtgag > ATTGGAaccac), Δ5′SS-GU (gt >cc), ΔBS (tgctgac > tgccgtt), ΔPPT (cctgtcccttttttttcc > ggagagggaaaaaaaagg), Δ3′SS (cagCT > accTC), Δ3′SS-AG (ag > cc), and ΔESE (TCA > AAT). Fourteen nucleotide long *Ron* ESE sequence is as reported before (42) (UGGCGGAGGAAGCA), *AdML* 5′SS RNA (UUGGGGUGAGUACU), and *β*-globin 5′SS (GGGCAGGUUGGUAU).

### *In vitro* reconstitution and purification of U1 snRNP

For reconstitution and purification of U1 snRNP, full-length U1 snRNA was transcribed in large scale *in vitro* using run off transcription from T7 promoter and treated with DNase I. U1 snRNP was assembled as described before (30) and purified by anion exchange chromatography (Mono Q; Cytiva) using a KCl gradient (from 250 mM KCl through 1M KCl). Particles were flash-frozen in liquid N_2_ and stored at −80 °C in single-use aliquots.

### *In vitro* assembly and purification of pre-mRNA complexes

*AdML* IVS1 was synthesized by run-off transcription and treated with DNase I (New England Biolabs). For chromatographic purification, reconstitution of pre-mRNA complexes was carried out in 500 μl volume except for where U2AF65 and U2AF35 were added; in the latter cases, reconstitution was carried out in 1 ml volume. 100 nM pre-mRNA in complex with 300 nM MBP-MS2 was incubated with 300 nM U1 snRNP for 5 min at 30 °C in 20 mM HEPES-NaOH pH 7.5, 250 mM NaCl, 2 mM MgCl_2_, 1 mM DTT, 0.5 M urea, 0.3% poly(vinyl alcohol) (PVA, Sigma P-8136), and 5% glycerol. Then SRSF1 (500 nM) was added where indicated and incubated for 5 min at 30 °C; thereafter, U2AF65 (150 nM), SF1_320_ (150 nM), and U2AF35 (150 nM) were added where indicated and incubated further for 5 min at 30 °C. Where SRSF5 (500 nM) is added instead of SRSF1, it is added before U1 snRNP. Then the reaction was incubated at room temperature for 5 min, followed by high-speed centrifugation at 4 °C for 2 min. The complexes were then bound to 1 ml Mono Q column (Cytiva) in 20 mM HEPES-NaOH pH 7.5, 250 mM NaCl, 2 mM MgCl_2_, 1 mM DTT at 4 °C. The elution buffer had identical components as the binding buffer except for having 2M NaCl instead of 250 mM. Complexes formed with *AdML* variants were eluted with a combinatorial gradient from 0 – 20% of the elution buffer for 5 ml followed by 20 – 30% for 10 ml. Each fraction size was 330 μl. The pre-mRNA-containing fractions were concentrated by binding to amylose resin (New England Biolabs). For SDS PAGE analysis, the resin beads were boiled in Laemmli sample buffer. For RNA isolation, the resin beads were incubated with proteinase K (New England Biolabs) in proteinase K buffer followed by phenol:chloroform extraction and ethanol precipitation. The precipitated RNA was directly dissolved in 9M urea dye for analysis by urea PAGE. We routinely observed a proportion of the pre-mRNA precipitated in the presence of U1 snRNP on the column and this proportion was slightly higher for the recruitment-defective pre-mRNAs.

For amylose pull-down assay (without chromatographic separation), the reaction was rotated with amylose resin for 20 min at room temperature followed by 5 min at 4 °C. Thereafter, the supernatant was removed by gentle centrifugation, the resin was washed with chilled binding buffer (same as binding buffer in chromatography experiments), followed by boiling in Laemmli sample buffer for SDS PAGE analysis.

### Negative stain electron microscopy

The EM grid was prepared by depositing particles on the Carbon-Formvar grid (01754-F F/C 400 mesh Cu from Ted Pella) activated by 30 seconds of glow discharge, briefly washing twice with deionized water, and staining with 0.5% Uranyl Acetate for 1 min. Images were collected in FEI Tecnai G2 Sphera. 2D-average analysis was carried out using EMAN2 program.

### *In vitro* selective 2′ hydroxyl acylation followed by primer extension (SHAPE) by mutational profiling (SHAPE-MaP)

To detect both stable and dynamic interactions between the pre-mRNA and U1 snRNP or SR proteins, we carried out SHAPE-MaP experiment with protein-free, U1 snRNP-bound, or SR protein-bound pre-mRNAs using 2 mM NMIA; low NMIA concentration was beneficial for detecting weak interactions. For carrying out SHAPE with 2 mM N-Methylisatoic anhydride (NMIA, Millipore-Sigma) with or without U1 snRNP, RNA was denatured in the presence of 5 mM EDTA pH 8.0, 50 mM NaCl at 95 °C for 3 min and then renatured by immediately placing on ice for 15 min. After neutralization of EDTA with MgCl_2_, pre-mRNAs were incubated at 30 °C under the same conditions used for assembly of the pre-mRNA complexes as indicated above in 1 ml volume. First, the MS2-tagged pre-mRNAs were bound to 3X MBP-MS2 protein, then either U1 snRNP or the U1 snRNP storage buffer (20 mM HEPES pH 7.5, 400 mM KCl, 2 mM MgCl_2_, 15 mM Arginine-HCl, 15 mM Glutamate-KOH, 5 mM DTT) or an SR protein or the SR protein storage buffer was added to the reaction mixture. After incubation, the complexes were mixed with amylose resin (New England Biolabs) and were rotated at 4 °C for 30 min. Next, the resin was separated from the supernatant by gentle centrifugation and complexes bound to the resin were eluted with 400 μl of the reaction buffer with 20 mM maltose by rotating the tubes for 1 hour at 4 °C. The pre-mRNA-containing solutions were then transferred to fresh tubes containing 6.67 μl 120 mM NMIA solution (in DMSO) or the same volume of DMSO and mixed by pipetting immediately. Then, the tubes were kept at 16 °C for 1 hour. Finally, the RNA was purified by RNeasy Mini Kit (Qiagen) including a proteinase K digestion step following manufacturer’s instructions.

For modeling secondary structure of the pre-mRNA variants, we carried out SHAPE-MaP with the recommended high level of NMIA (8 mM), which was beneficial for generating high mutation rates. This might provide less information regarding the weak/transient interactions but differentiates non-transient/robust interactions of different strengths, thus aiding in generation of the thermodynamically stable secondary structural models. For carrying out SHAPE with 8 mM NMIA, the pre-mRNAs were not bound to any protein and the reaction buffer was same as with pre-mRNA complex formation. Denatured and renatured pre-mRNAs were incubated at 30 °C in the reaction buffer for 5 min in 1 ml volume at 100 nM concentration, then the reactions were transferred to fresh tubes containing 66.7 μl 120 mM NMIA in DMSO or DMSO alone and incubated at 30 °C for 30 min. Then RNA was precipitated from the reaction with 300 mM sodium acetate and 3X volume of ethanol and the pellet was dissolved in 50 μl water. Then the RNA was purified by Monarch RNA cleanup kit (NEB).

The SHAPE reactivity of protein-free pre-mRNAs obtained by the two methods described above are likely to differ at places primarily because of differential reactivity of individual nucleotides at different concentrations of NMIA. Additionally, RNA passed through a long processing step before treatment with NMIA as in the first case is likely to cause alteration in certain structural features of the RNA.

Denatured control for each RNA was generated as previously described before (43) where denatured RNA was treated with 2 mM or 8 mM NMIA for 5 min at 95 °C.

Reverse transcription was carried out in MaP buffer as described before (43) and sequencing library was prepared by limited cycle PCR amplification. The samples were deep-sequenced in multiplex using MiSeq platform (Illumina) using MiSeq Nano kit (2x 250 bp, paired end) at the IGM Genomics Center, University of California, San Diego, La Jolla, CA. The demultiplexed FastQ Files obtained from the facility were processed using ShapeMapper 2.0 pipeline (43) following authors’ instructions to obtain the SHAPE reactivity. All negative SHAPE reactivity values were considered zero.

### DREEM analysis of SHAPE-MaP data of RNAs treated with 8 mM NMIA

<u>D</u>etection of <u>R</u>NA folding <u>e</u>nsembles using <u>e</u>xpectation-<u>m</u>aximization (DREEM) (44) analysis reveals alternative thermodynamically stable conformations assumed by an RNA molecule from MaP-Seq data. It segregates the reactivity profile into two classes based on the incidence of co-occurrence (or the lack of it) of mutations in a single RNA molecule. Since the authors of the script used DREEM to analyze DMS modification, which modifies only A and C, the script is written to include A and C only. We changed the Python Boolean in line 111 in the main script to include T and G, since NMIA modifies all four nucleotides. The pipeline runs the expectation-maximization algorithm 10 times generating 10 possible combinations of two classes, referred to as K2 classes (run_1 through run_10). DREEM also runs to club together all mutations into one reactivity profile, referred to as the K1 class (run_1). Then the models are tested using corresponding Bayesian information criteria (BIC) values of the K1 run and each of the K2 runs. A lower BIC value of a K2 run compared to that of the K1 run indicates that segregation of the SHAPE reactivity into two classes in that K2 run is statistically feasible.

### Transfection-based splicing assay

Splicing assay was carried out as described before (18). Antisense morpholino oligonucleotide (AMO) for the 25-nt region at the 5 end of U1 snRNA (U1 AMO) was as described before (45): 5′-GGTATCTCCCCTGCCAGGTAAGTAT-3′, scrambled AMO: CCTCTTACCTCAGTTACAATTTATA.

### *In vivo* SHAPE

*HeLa* cells grown in a six-well plate up to ~ 80% confluency were transfected with 5 nmol U1 AMO and after 12 hours were transfected with pre-mRNA constructs. After six hours, cells were treated with 100 mM 2-methylnicotinic acid imidazolide (NAI, Sigma) (46) for 15 minutes at 37 °C followed by quenching of the SHAPE reagent with 125 mM 1,4-dithiothreitol (DTT). Isolation of total RNA and DNase I treatment were carried out as with *in vivo* splicing assay (18). Denatured control was prepared as described before (43) by treating the total RNA with 100 mM NAI at 95 °C for 5 min. SHAPE-MaP reaction, deep-sequencing, and data analysis were carried out as described above. The per-nucleotide mapped read depth for all *in vivo* samples were maintained at 30000 or higher (instead of the recommended depth of 2000 (47)) to include the possible biological variabilities in *in vivo* SHAPE reactivity of each nucleotide.

### Two-color fluorescence measurement

RNA was denatured in 50 mM NaCl and 5 mM EDTA pH 8.0 at 95 °C for 3 min and then placed on ice. Then 200 nM RNA was mixed with 200 nM DNA probes, heated at 55 °C for 5 min, and then gradually cooled to room temperature over a period of 60 min. This allowed the DNA probes to anneal to the loop areas of the folded RNA. Three samples were prepared for each RNA: one containing Cy3-labeled 5 probe and Cy5-labeled 3′ probe, one containing Cy3-labeled 5′ probe and unlabeled 3′ probe, and one containing unlabeled 5 probe and Cy5-labeled 3′ probe. Next, EDTA was neutralized with MgCl_2_. Thereafter, HEPES pH 7.5, NaCl, MgCl_2_, and PVA were added to the final concentration of 20 mM, 250 mM, 2 mM, and 0.3%, respectively, and the reaction was incubated at 30 °C for 5 min. Fluorescence was measured in Jasco Spectrofluorometer 8500 with a bandwidth of 5 nm for excitation and emission. All measurements with intact *AdML* were replicated thrice from distinct samples. For annealing of the structure-disrupting 40-nt long DNA to *AdML* RNA, 200 nM RNA, 200 nM disruptor DNA, and 200 nM labeled DNAs were mixed in the presence of 50 mM NaCl and 5 mM EDTA pH 8.0 and heated at 95 °C for 3 min. Thereafter, the mixture was gradually cooled for over 60 min to room temperature.

### Electrophoretic mobility shift assay (EMSA)

EMSA was carried out as described before (18). Six-percent (80:1) polyacrylamide gel was used for EMSA with 14-nt long RNA, which was purchased from Integrated DNA Technologies and end-labeled with T4 polynucleotide kinase (NEB) and γP^32^-ATP (6000 Ci/mmol; 10 μCi/μl) following manufacturer’s instructions.

### GST pull-down assay

GST pull-down assay was carried out as described before (31).

## RESULTS

### *AdML* splice signal mutants with intact 5′SS are defective in U1 snRNP recruitment

For testing functional U1 snRNP binding to *AdML*, purified recombinant U1 snRNP particle (see Supplementary Figures S1A-S1C) was mixed with WT *AdML* (in chimera with 3X MS2-protein binding loops) and MBP-MS2 fusion-protein as described before (48), and the resulting complexes were purified by anion-exchange chromatography. The chromatogram (Figure 1A – blue line) revealed one distinct and two overlapping peaks. SDS (Figure 1B) and urea (Supplementary Figure S1D) PAGE analyses of the peak fractions showing protein and RNA, respectively, indicate that the first peak (fractions 19-21) contains free U1 snRNP, the second peak (fractions 24-29) contains the U1 snRNP:pre-mRNA complex (MBP-MS2 represents pre-mRNA but at 3 times the pre-mRNA level), and the third peak (fractions 30-32) contains diminishing levels of U1 snRNP and high levels of MBP-MS2, suggesting trailing of the U1 snRNP: pre-mRNA complex overlapping with excess free pre-mRNA. The fractions indicated as “pre-mRNA complexes” (fractions 24-32 in Figure 1B) were concentrated by MBP-MS2-pull-down using amylose resin before PAGE analysis.

**Figure 1.**
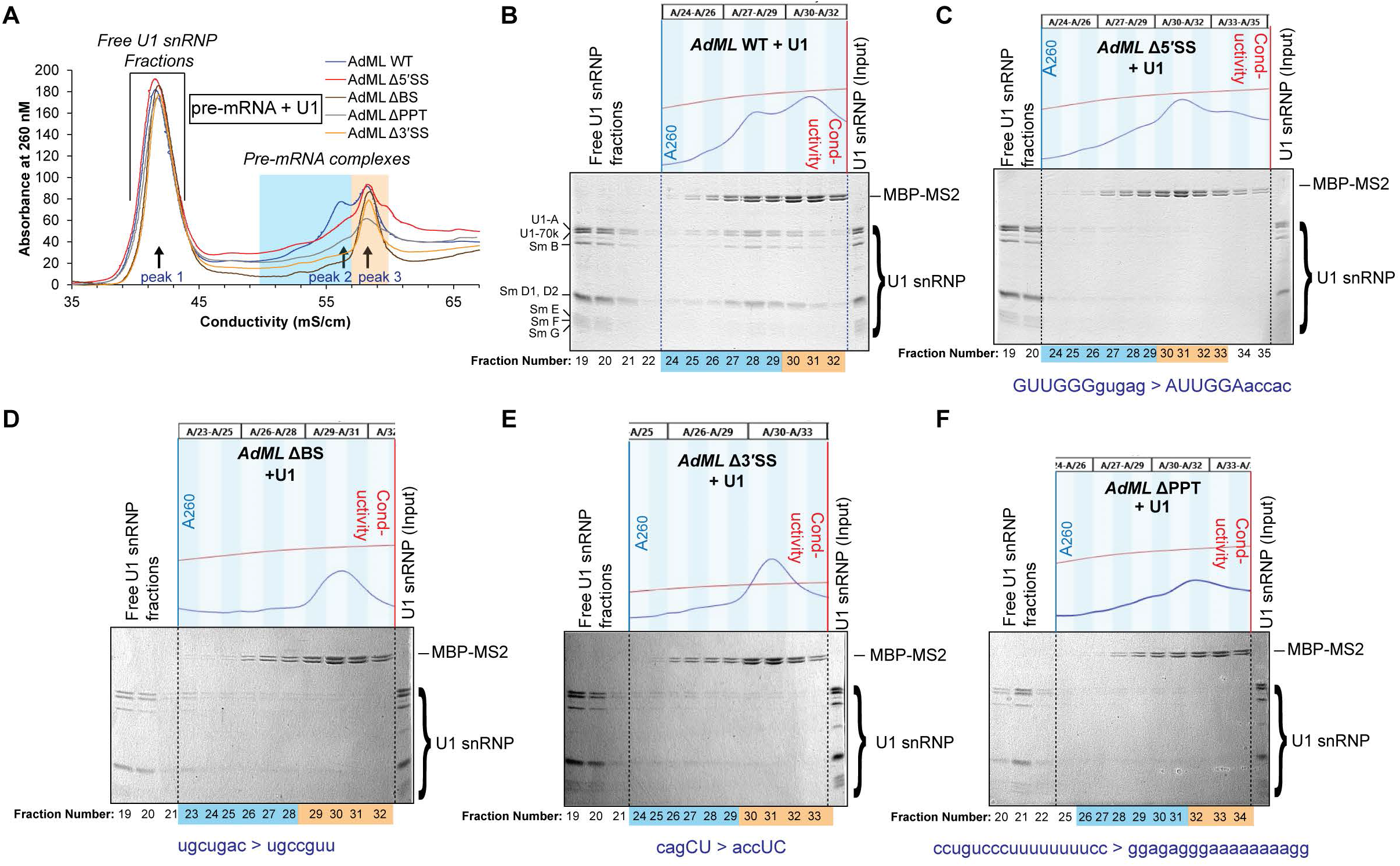
*AdML* splice signal mutants with intact 5′SS are defective in U1 snRNP recruitment. (A) Chromatograms of purification of U1 snRNP:pre-mRNA:3XMBP-MS2 complexes by anion-exchange chromatography with an NaCl gradient; conductivity instead of volume is plotted along *x*-axis to reduce shifts in peak positions caused by inconsistency in salt gradient generation by FPLC; peaks 1, 2, 3 of WT *AdML* chromatogram (a clear peak 2 is absent in the chromatograms of *AdML* mutants) are indicated with arrows; each of the fractions within the region indicated to contain ‘pre-mRNA complexes’ were concentrated by pull-down with amylose resin for analysis by Coomassie-stained SDS PAGE; blue shade corresponds to peak 2 and preceding fractions of *AdML* WT complexes and equivalent fractions of *AdML* mutant complexes; orange shade corresponds to fractions within peak 3. (B, C, D, E, F) SDS PAGE of *AdML* WT (B), *AdML* Δ3′SS (E) and *AdML* ΔPPT (F) fractions corresponding to the chromatogram shown in (A); 2.5 pmol U1 snRNP was used in the ‘input’ lane in the SDS gel; blue and orange shade shown at the bottom of the gel images corresponds to fractions within the region of the chromatogram shaded with the same color; fraction-numbers of the analyzed samples are provided under each lane; mutated sequence for *AdML* mutants is also indicated under the SDS gel images; the raw chromatogram representing the fractions containing the pre-mRNA complexes are shown above each gel, where blue line (absorbance trace) and red line (conductivity trace) are placed along primary and secondary *y-*axes, respectively.

Next, we examined U1 snRNP binding to the splice signal mutants of *AdML*. Similar experiments with 3X MS2-tagged *AdML* mutants (Δ5′SS, ΔBS, Δ3′SS, ΔPPT) (Figure 1A) indicate that the level of U1 snRNP co-eluted with *AdML* mutants is distinctively less than that for the WT substrate (Figures 1C, 1D, 1E, 1F). A representative urea gel demonstrating the total RNA content (pre-mRNA and U1 snRNA) in the fractions is also shown for *AdML* Δ5′SS (Supplementary Figure S1E) suggesting significantly weaker co-elution of U1 snRNA with *AdML* Δ5′SS compared to *AdML* WT. The chromatograms for *AdML* Δ3′SS and ΔBS exhibited abolition of the second peak (Figure 1A) while those for *AdML* Δ5′SS and ΔPPT exhibited a shoulder in place of the second peak. The shoulder likely represents certain conformational state(s) of the mutant pre-mRNAs.

The importance of 5′ and 3′ exonic segments in U1 snRNP recruitment was also examined by the chromatographic U1 snRNP binding assay. Earlier literature indicates that loops immediately upstream of the 5′SS could be important for early spliceosome assembly (18). We assessed involvement of these loops in recruiting U1 snRNP by examining binding of U1 snRNP to the splicing-defective exonic hybridization (EH) mutant (18). This mutant contains several altered nucleotides (spanning positions 29 to 55, as numbered in Figure 2A) upstream of the 5′SS (18) to eliminate loops in this region. Comparative RNA profile analysis in chromatographic fractions (Supplementary Figure S1F) of WT vs *AdML* mutant complexes showed a markedly reduced co-elution of U1 snRNA with the *AdML* EH mutant (Supplementary Figure S1G) suggesting their weak interaction. We also performed a similar analysis with an *AdML* construct lacking the 3′ exon (*AdML* Δ Ex2 (Supplementary Figure S1H) was also distinctively low (Supplementary Figure S1D).

**Figure 2.**
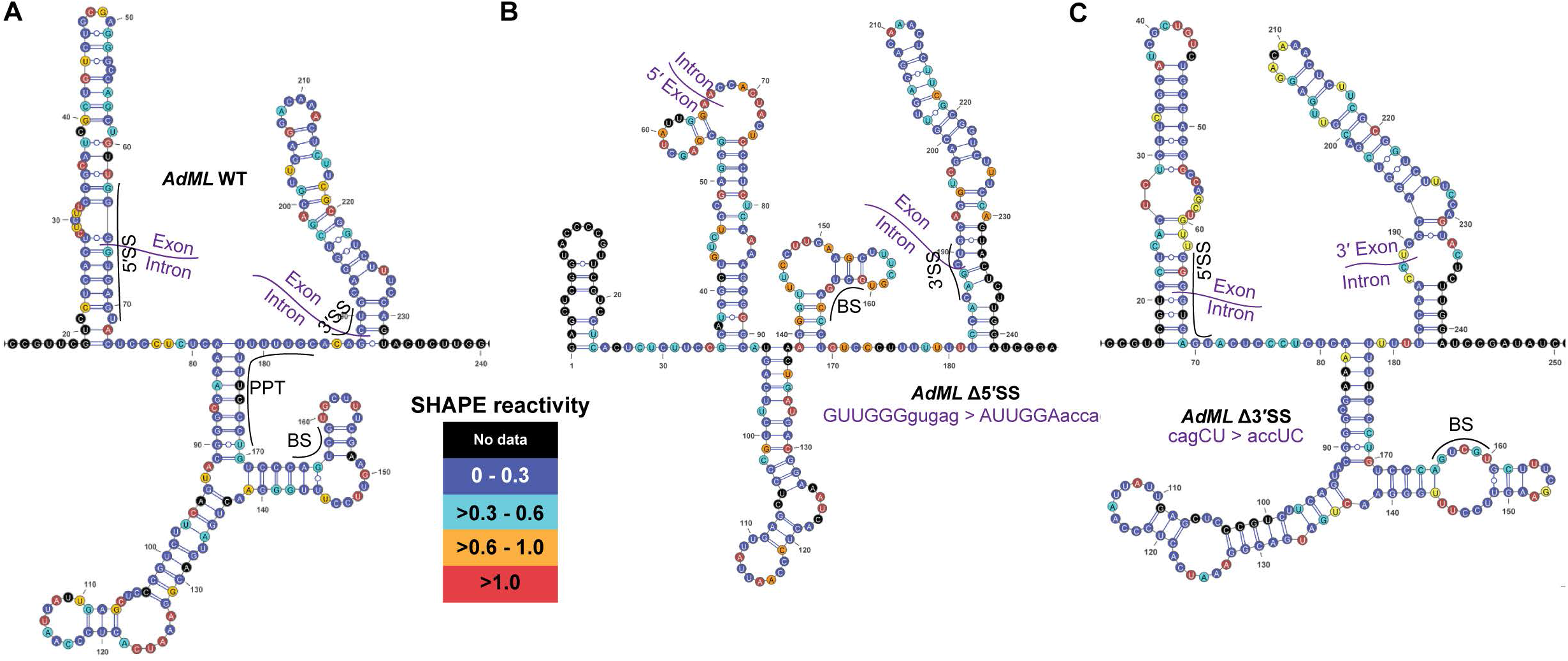
Mutation in major splice signals disrupts global structure of *AdML*. (A, B, & C) SHAPE-derived secondary structure models of *AdML* WT (A), *AdML* Δ5′SS (B), and *AdML* Δ3′SS (C); nucleotides are color-coded according to their SHAPE reactivity (see associated legend); positions of splice signals are indicated; position of PPT is shown only with the WT substrate for clarity; the mutated sequence is shown with each model.

We then examined U1 snRNP binding to radiolabeled 14-nt long *AdML* 5′SS RNA by EMSA. No binding to U1 snRNP, even at 400 nM concentration, was detected (Supplementary Figure S1I – lanes 2, 3, 4). In contrast, U1 snRNP binding to radiolabeled full-length *AdML* appears to be efficient at 160 nM concentration (Figure 5A, lane 6). As a positive control, we used SRSF1-RBD (Supplementary Figure S5A), which efficiently binds to sequences containing GGN and CN (N=any nucleotide) (49), also found in *AdML* 5′SS RNA.

These data suggest an important role of the global structural landscape of *AdML* pre-mRNA in U1 snRNP recruitment.

### Mutations in major splice signals disrupt the global three-dimensional structure of *AdML*

To understand why the splice signal mutants exhibited a weaker U1 snRNP recruitment efficiency compared to the WT substrate even when 5′SS was intact, we hypothesized that mutation in the splice signal sequences disrupts the three-dimensional structural scaffold of *AdML*. Since a difference in SHAPE reactivities of the WT and the mutants would be reflective of their different/altered three-dimensional structures (secondary and tertiary), we measured SHAPE reactivity of *AdML* WT, Δ5′SS, Δ3′SS, ΔBS using the SHAPE-MaP technique (43) with 8 mM N-Methylisatoic anhydride (NMIA) and derived their secondary structure models with RNAstructure (50). Each of the *AdML* splice signal mutants exhibited a different SHAPE reactivity profile and a distinct secondary structure model compared to those of the WT substrate (Figure 2A) – *AdML* Δ5′SS (Figure 2B), *AdML* Δ3′SS (Figure 2C), and *AdML* ΔBS (Supplementary Figure S2A). The differences in the secondary structure models also conformed to differences in their tertiary structures. The raw SHAPE reactivity values are provided in Supplementary File 1.

Since several nucleotides within each splice signal were mutated, the resulting structural disruption of the pre-mRNA might not be reflective of splice signals holding key positions in the global structural scaffold of *AdML* pre-mRNA. Accordingly, we mutated just the conserved nucleotides of 5′- and 3′-SS (i.e GU and AG) generating *AdML* Δ5′SS-GU and *AdML* Δ3′SS-AG, respectively. SHAPE reactivity and the derived secondary structural models of these mutants were also different from that of WT *AdML* (Supplementary Figure S2B, S2C, Supplementary File 1). We also tested the significance of this structural disruption in the *AdML* Δ3′SS-AG mutant by examining U1 snRNP recruitment by chromatography (Supplementary Figure S2D) and observed a weaker coelution of U1 snRNP with the mutant pre-mRNA (Supplementary Figure S2E).

We then carried out DREEM analyses (44) of the MaP-Seq data of NMIA-treated *AdML* mutants to examine the possibility of alternative thermodynamically stable conformations (Methods). The Bayesian information criteria (BIC) value of none of the ten K2 runs was lower than that of the K1 run for any of the *AdML* variants (Supplementary Figure S2F). This suggests that the proportion of RNA molecules assuming additional thermodynamically stable structure(s) is either zero or too small to be detected.

Overall, this result suggests that the major splice signals are functionally and structurally integrated into the global structural scaffold of *AdML* pre-mRNA.

### Strandedness of nucleotides across the pre-mRNA correlates to U1 snRNP recruitment

We hypothesized that in order for the global three-dimensional structural scaffold of *AdML* to regulate U1 snRNP recruitment, U1 snRNP could potentially contact various nucleotides across the pre-mRNA, which are spatially proximal to the 5′SS region. Therefore, we examined potential points of contact between U1 snRNP and *AdML* by SHAPE experiments with 2 mM NMIA (see Methods), followed by calculating the SHAPE reactivity differential of U1 snRNP-bound and protein-free (mock-treated) *AdML* ([*AdML*+U1] – *AdML*) (calculated by subtracting SHAPE reactivity of each nucleotide of *AdML* WT from that of the corresponding nucleotide of *AdML* in complex with U1 snRNP) (blue line in Figure 3A as well as 3B) (raw SHAPE values provided in Supplementary File 1 and plotted in Supplementary Figure S3A). A lower SHAPE reactivity of a nucleotide of U1 snRNP-bound *AdML* compared to that of protein-free WT *AdML* indicates its protection, caused by direct contact with U1 snRNP components or base-pairing introduced by pre-mRNA structural remodeling. On the other hand, a higher SHAPE reactivity within U1 snRNP-bound *AdML* means removal of structural constraints through pre-mRNA structural remodeling. We observed changes in SHAPE reactivity across the entire pre-mRNA upon U1 snRNP binding to *AdML*: distinct enhancement of flexibility was observed upstream of the 5′SS, in the middle of the intron at a terminal loop area (110-127 nt), around the BS region, immediately upstream of the 3′SS, and at two places in the 3′ exon (Figure 3A); enhanced protection was observed at various places in the 5′ exon, at and immediately downstream of the 5′ SS, slightly upstream of the BS (around nt 140), and immediately downstream of the 3′SS (Figure 3A).

**Figure 3.**
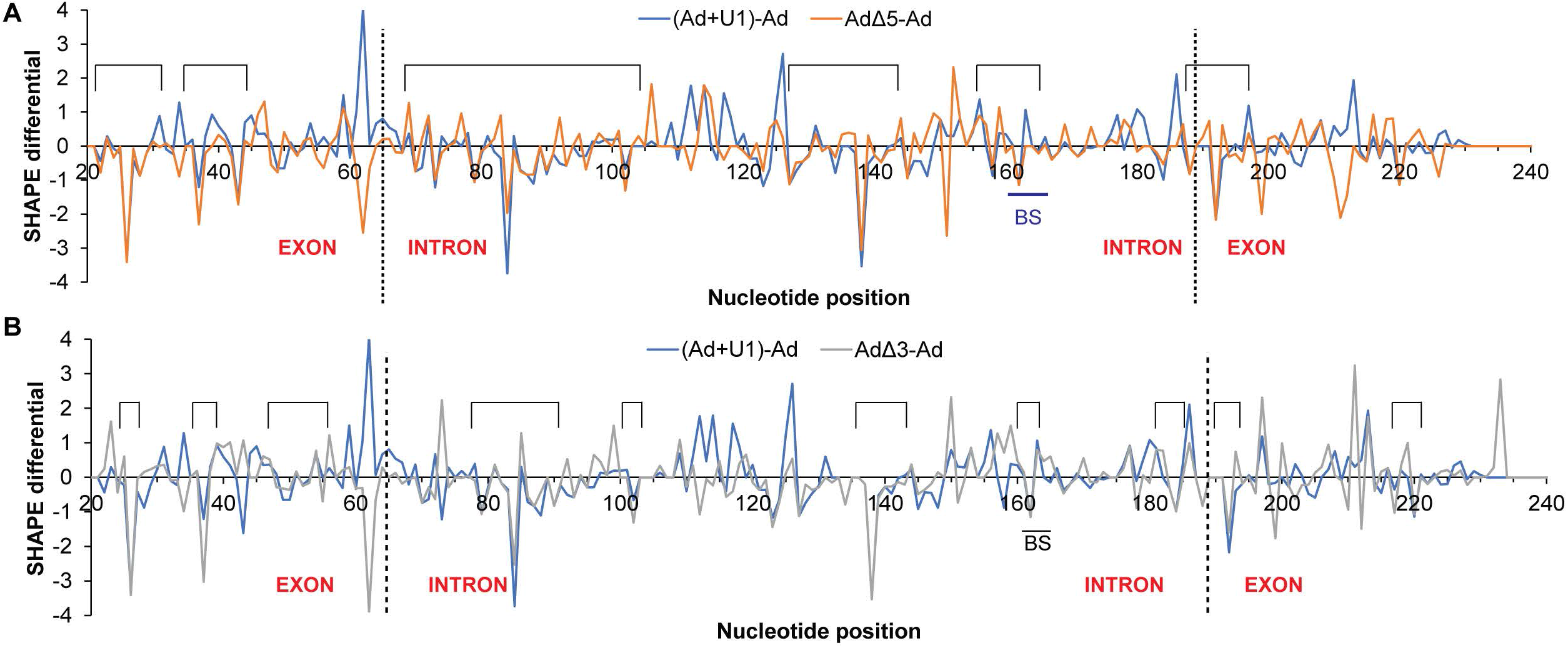
Strandedness of nucleotides across the pre-mRNA correlates to U1 snRNP recruitment. (A) SHAPE differentials [(Ad+U1) – Ad] (calculated by subtracting the SHAPE reactivity of each nucleotide of mock-treated *AdML* from that of the corresponding nucleotide of *AdML* in complex with U1 snRNP) and [Ad Δ5 – Ad] (calculated by subtracting the SHAPE reactivity of each nucleotide of mock-treated *AdML* from that of the mock-treated *AdML* Δ5′SS are overlaid onto each other; nucleotide positions exhibiting distinct negative values in both curves are indicated with rectangular bracket above the plot (these nucleotides loose SHAPE reactivity upon engagement of U1 snRNP potentially due to contact with U1 snRNP in *AdML* WT and are occluded by base-pairing in the protein-free *AdML* Δ5′SS); exon-intron junctions are indicated with dotted vertical lines. (B) SHAPE reactivity differentials of WT *AdML*+U1 snRNP complex and protein-free *AdML* i.e. [(Ad+U1) – Ad] and *AdML* Δ3′SS and *AdML* WT i.e. [Ad Δ3 – Ad] are overlaid onto each other.

Since splicing factors are reported to generally interact with single-stranded RNA segments (11,18), we examined if the *AdML* mutants that fail to bind U1 snRNP efficiently have some of the potential U1 snRNP contact points occluded by base-pairing. We estimated the SHAPE reactivity of protein-free *AdML* Δ5′SS and *AdML* Δ3′SS treated with 2 mM NMIA (raw SHAPE data shown in Supplementary File 1 and plotted in Supplementary Figures S3B and S3C). Then we calculated the SHAPE reactivity differentials (*AdML* Δ5′SS – *AdML* WT) (orange line of Figure 3A) and (*AdML* Δ3′SS – *AdML* WT) (grey line of Figure 3B) by subtracting the SHAPE reactivity of each nucleotide of *AdML* WT from that of the corresponding nucleotide of *AdML* Δ5′SS or *AdML* Δ3′SS, respectively. Thereafter, we overlaid these differentials onto the plot of ([*AdML*+U1] – *AdML* WT) differential (blue line in Figures 3A and 3B). Interestingly, both (*AdML* Δ5′SS – *AdML* WT) and (*AdML* Δ3′SS – *AdML* WT) differentials exhibited low negative values at several common nucleotide positions with ([*AdML*+U1] – *AdML* WT), marked with rectangular brackets above the plots.

Overall, these results suggest that availability of certain single-stranded nucleotides across the pre-mRNA correlates to U1 snRNP recruitment efficiency to *AdML* variants.

### Close spatial positioning of splice signals to 5′SS

Many nucleotides across the pre-mRNA exhibit a reduction in SHAPE reactivities upon U1 snRNP engagement, some of which potentially contact U1 snRNP components. To accommodate this, we propose that these nucleotides will have to be spatially close to each other and in proximity to the 5′SS. Accordingly, we examined if the pre-mRNA helices could come close to each other in three-dimensional space in the protein-free state by Förster resonance energy transfer (FRET). We annealed a Cy3-labeled short DNA (20-nt long DNA #2) to the terminal loop area in the 5′ exon (around nt-50 of helix A) and a Cy5-labeled short DNA to either the intronic stem-loop around nt-115 of helix B (20-nt long DNA #5), the intronic stem-loop around nt-155 of helix C (19-nt long DNA #3), or the 3′ exonic stem-loop around nt-210 of helix D (16-nt long DNA #6) (Figure 4A) to *AdML*. To avoid major disruption of global structure of the pre-mRNA, *AdML* was first refolded and then the primers were annealed (see Materials and Methods). Since disruption of the base-pairing immediately upstream of the 5′SS is important for splicing (18), perturbation of certain base-pairs in this region due to annealing of the Cy3-labeled DNA #2 is not likely to affect the relevant pre-mRNA structure. For the other terminal loops, the labeled DNAs anneal to largely single-stranded regions with minimal perturbation of local structure. Steady-state fluorescence was monitored by exciting at 550 nm and scanning for fluorescence-emission intensity at wavelengths between 560 and 720 nm (Supplementary Figures S4A, S4B, S4C). Cy3 emission (maximum at 565 nm) was reduced in the presence of the Cy5 probe with all three pairs of DNA probes (2+3, 2+5, 2+6) suggesting FRET. A corresponding gain of Cy5 emission (emission maximum 665 nm) was also noted in the presence of Cy3. The level of reduction in Cy3 emission upon addition of the Cy5 probe was significantly higher with the DNA probe-pairs 2-3 and 2-6 than 2-5 (Figure 4B) suggesting that annealed DNA #2 (helix A) is further from annealed DNA #5 (helix B) than annealed DNAs #3 and #6 (helices C & D). This is reflective of early-stage proximity of 5′SS (located in helix A) with BS (in helix C) and 3′ (in helix D) in protein-free *AdML*.

**Figure 4.**
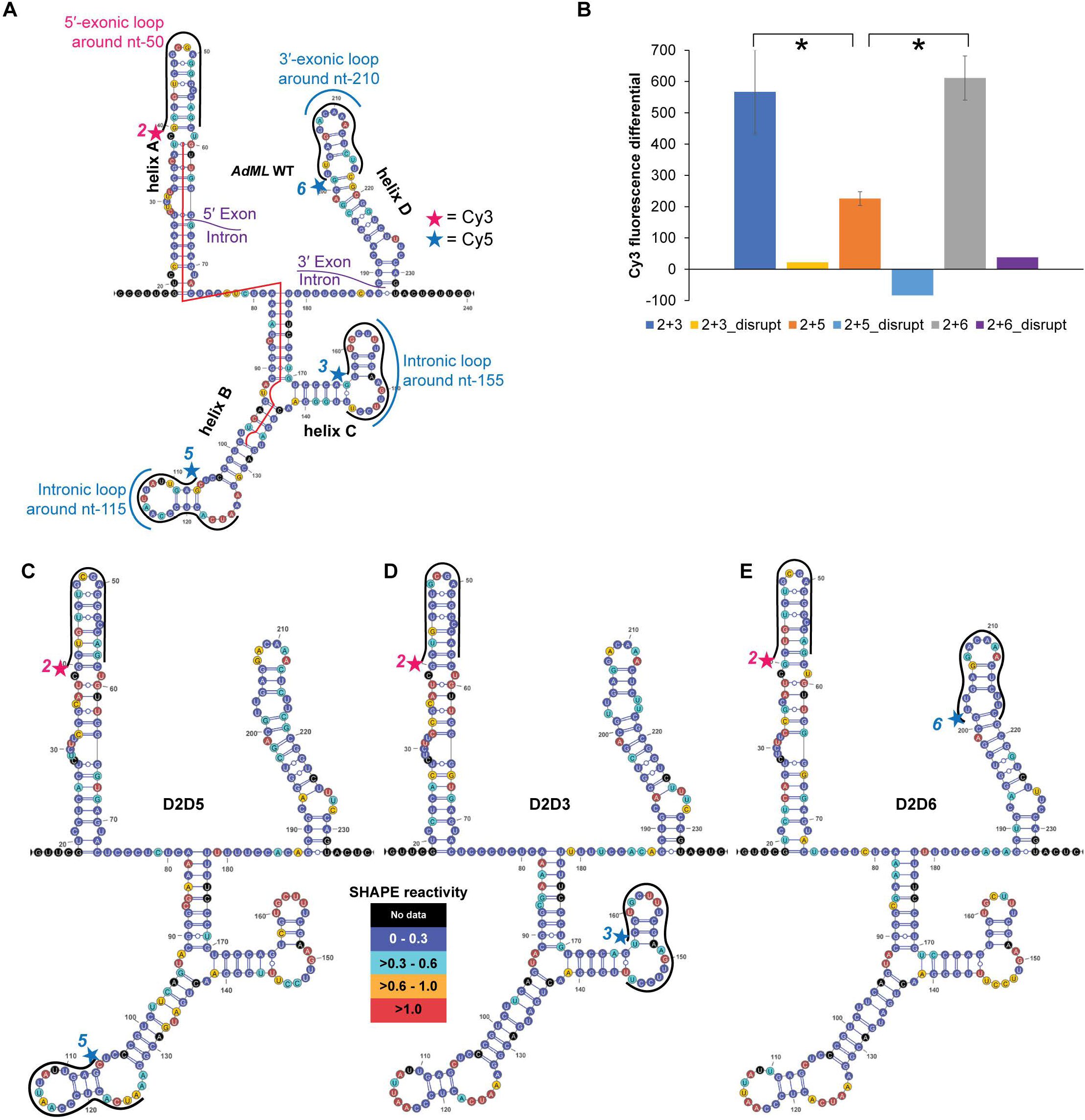
Close spatial positioning of splice signals to 5′SS. (A) SHAPE-derived secondary structure models of WT *AdML* showing the positions of the Cy3- and Cy5-labeled DNA probes; Cy3-labeled DNA probe is labeled as 2 and the remaining probes as 5, 3, and 6; the position of the 40-nt long structure-disrupting DNA is shown as a red line; the four helices of *AdML* are labeled as A, B, C, D. (B) Reduction in Cy3 emission upon hybridization of Cy5 probe to protein-free *AdML* WT shown for probe-pairs 2-3, 2-5, and 2-6; error bars indicate standard deviation (n=3); ‘*’ indicates statistical significance (*p* < 0.05) obtained by one-tailed t-test; the same differential is also shown for *AdML* WT annealed to the disruptor DNA (n=1). (C, D, E) Secondary structure model of protein-free *AdML* WT with SHAPE reactivities of *AdML* annealed to the fluorescent DNA #2 and #5 (C), #2 and #3 (D), and #2 and #6 (E) overlaid onto it by color-coding of the nucleotides as per the associated legend.

We then measured the SHAPE reactivities of *AdML* WT pre-mRNA annealed to these pairs of DNA probes with 8 mM NMIA, which were then overlaid onto the secondary structure model of WT *AdML* (Figure 4C, D, E). Loss of SHAPE reactivity upon hybridization of the DNA probes was observed in all cases but to different extents. In addition, the intensity of SHAPE reactivity in all models correlated well with the strandedness of protein-free WT *AdML*. Overall, these results suggest that a major structural perturbation was not introduced upon annealing of the DNA probes.

We also examined effects on FRET with all three probe pairs upon disruption of *AdML* global structure. We disrupted the structure by annealing with a long DNA (red line in Figure 4A) or by reducing the Mg^2+^ concentration from 2 mM to 0.1 mM, since Mg^2+^ ions are important for the RNA tertiary structure (51). We observed no significant FRET with any of the probe pairs upon disruption with the disruptor DNA (Figure 4B, Supplementary Figure S4D, S4E, S4F). Additionally, under the low Mg^2+^ condition, FRET values obtained with all three pairs of probes were greatly diminished (Supplementary Figure S4G, S4H, S4I). The raw fluorescence values are provided in Supplementary File 2.

These results suggest that splice signals residing in different helices of *AdML* could come close to each other in the protein-free state for recruiting the early spliceosomal components.

### The ESE-dependent RBP SRSF1 stabilizes U1 snRNP on WT *AdML* but not *AdML* Δ3′SS

For mammalian pre-mRNAs, functional recruitment of the early spliceosomal components at the splice signals are reported to require interactions with ESE-bound RBPs. Since we observed stable and specific U1 snRNP recruitment to protein-free *AdML*, we hypothesized that the ESE-dependent functionalities of RBPs could stabilize U1 snRNP to further prepare the complex for the next step of the assembly. Therefore, we tested the stability of the U1 snRNP:*AdML* complex in the presence as well as the absence of ESE-dependent SR protein SRSF1 – known to stabilize U1 snRNP on the pre-mRNA through direct interaction (24,31,52) – by EMSA. Supplementary Figure S5A shows the SR protein constructs used in this study, all of which are reported to be functional. The truncated SRSF1 containing only the RNA binding domain (RBD) is referred to as SRSF1-RBD. SRSF1 is activated upon release of its RBD from the C-terminal Arg-Ser-rich (RS) domain upon phosphorylation of the latter (31). In addition, biochemical and structural properties of the RBD of SRSF1 are sufficient to explain its sequence-specific binding, intermolecular interactions, and removal of base-pairing constraints (49). Functionally, the RBD of SRSF1 is active in splicing (53), inducing structural remodeling of the pre-mRNA (18), promoting assembly of the early spliceosome (18), and interacting with U1 snRNP components (31). Accordingly, we used the RBD to study the ability of SRSF1 to stabilize U1 snRNP on *AdML* pre-mRNA. *AdML* formed complexes with U1 snRNP both in the presence and the absence of SRSF1-RBD (Figure 5A – lanes 3 & 6). The *AdML*:U1 snRNP complex withstood the challenge of an excess of unlabeled 14-nt long *AdML* 5′SS RNA in the presence of SRSF1 better than in the absence of SRSF1 as indicated by the release of free radiolabeled probe upon the challenge (compare lanes 4 and 7). Challenging the *AdML* complexes with 14-nt long 5′SS of β-*globin* pre-mRNA (54) released less free probe than *AdML* 5′SS RNA (compare lanes 7 & 8). Complexes formed with *AdML* Δ5′SS pre-mRNA were significantly reduced in all cases (lanes 11-16) indicating a weaker interaction. Then we tested if the presence of SRSF1 could enhance U1 snRNP binding to one of the *AdML* mutants with intact 5′SS. Chromatographic purification of the mixture of *AdML* Δ3′SS, SRSF1, and U1 snRNP (Figure 5B), and subsequent SDS PAGE analysis of the fractions containing the pre-mRNA complexes (Figure 5C) suggest that U1 snRNP binding to *AdML* Δ3′SS was not enhanced to the extent of its binding to *AdML* WT (compare Figures 1B, 1E, and 5C).

**Figure 5.**
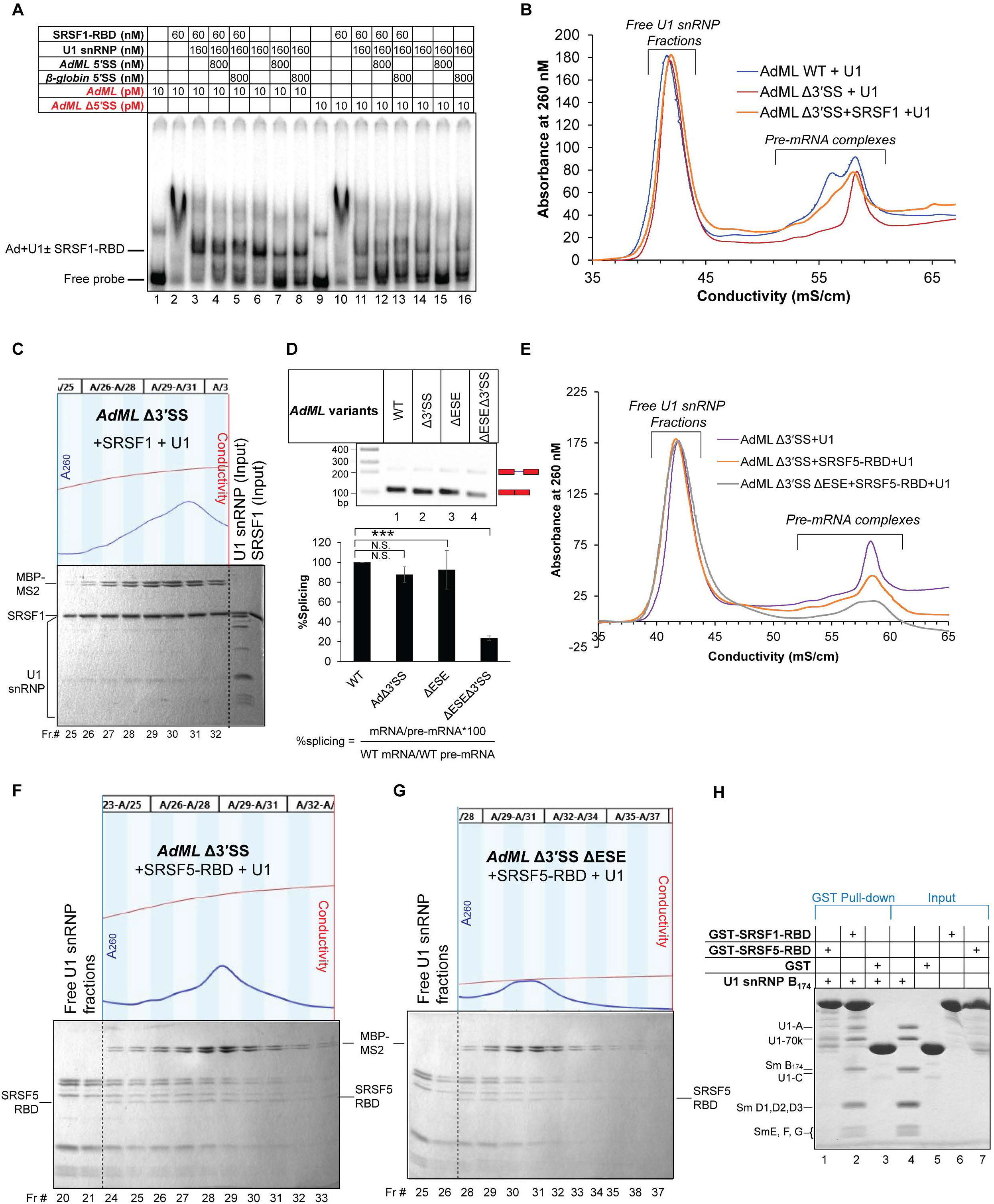
Differential functionalities of SR proteins with *AdML* WT and *AdML* Δ3′SS. (A) Electrophoretic mobility shift assay comparing the stability of U1 snRNP:*AdML* complex formed in the presence (lanes 3, 4, 5) or the absence (lanes 6, 7, 8) of SRSF1-RBD by challenging the complexes with short 14-nt 5′SS RNA to visualize the level of released free radiolabeled probe; all U1 snRNP-dependent complexes formed more weakly with *AdML* Δ5′SS (lanes 9-16); red script indicates radioactive components in the gel. (B) Chromatogram of purification of U1 snRNP-dependent complexes formed with *AdML* Δ SS in the presence of SRSF1; the chromatograms of purification U1 snRNP-dependent complexes formed with protein-free *AdML* WT and protein-free *AdML* Δ3′SS are shown for comparison. (C) SDS PAGE analysis of the pre-mRNA complexes formed with U1 snRNP and *AdML* Δ3′SS in the presence of SRSF1; full-length phosphomimetic SRSF1 stains five times more intensely than U1 snRNP proteins U1-A and U1-70k (Supplementary Figure S7K). (D) (Top) Transfection-based splicing assay of *AdML* WT and Δ3′SS with or without ESE mutation (mutated region shown in Supplementary Figure S5E); position of pre-mRNAs and mRNAs in the gel is indicated; (bottom) %splicing calculated from densitometric estimation of mRNA and pre-mRNA bands using the indicated formula of three biological replicas; error bar indicates standard deviation; N.S. = not significant, *** = *p* < 0.005 (one tailed one sample t-test). (E) Chromatograms of U1 snRNP-dependent complexes formed with *AdML* Δ3′SS and *AdML* Δ3′SS ΔESE in the presence of SRSF5-RBD; chromatogram of purification of mixture of U1 snRNP and protein-free *AdML* Δ3′SS is shown for comparison. (F, G) SDS PAGE of fractions corresponding to the pre-mRNA complexes formed with *AdML* Δ SS in the presence of U1 snRNP and SRSF5-RBD (F), with *AdML* Δ3′SS ΔESE in the presence of U1 snRNP and SRSF5-RBD (G); the raw SDS gel images are provided in Supplementary File 3. (H) GST-pull down assay showing absence of detectable interaction between U1 snRNP and GST-SRSF5-RBD (lane 1) and the presence of interaction between U1 snRNP and GST-SRSF1-RBD (lane 2); Sm B_174_ indicates Sm B (1-174 a.a.).

Overall, these results suggest that the pre-mRNA global structure dictates the functionality of the ESE-dependent RBP SRSF1 in stabilizing early spliceosomal components on the pre-mRNA.

### The RNA binding domain of SRSF5 promotes recruitment of U1 snRNP to *AdML* Δ3′SS

Although *AdML* Δ′SS could not recruit U1 snRNP in the protein-free state or in the presence of SRSF1, a transfection-based splicing assay indicated that *AdML* Δ3′SS splices as efficiently as *AdML* WT (Figure 5D – lanes 1 & 2) using a cryptic 3′SS (AG) 6-nucleotide downstream of the authentic 3′SS (Supplementary Figure S5B). Among the other splice signal mutants, *AdML* Δ5′SS and *AdML* ΔPPT demonstrated no splicing activity, while *AdML* ΔBS only weakly spliced into WT mRNA (determined by Sanger sequencing of the mRNA), likely using a different branch-site (Supplementary Figure S5C) (55). We also examined if U1 snRNP is required for splicing of *AdML* Δ3′SS, since U1 snRNP-independent splicing has also been reported (56). We expressed these variants of *AdML* in *HeLa* cells co-transfected with the 25-nt long antisense morpholino oligonucleotide complementary to the 25-nt of the 5′ end of U1 snRNA, as described previously (45). Inhibition of U1 snRNP caused accumulation of the pre-mRNA variants (Supplementary Figure S5D – compare lanes 1, 2 with 6, 7), which otherwise were quickly spliced into mRNAs.

The defect in U1 snRNP recruitment to *AdML* Δ3′SS could largely be attributed to an unfavorable global three-dimensional structural scaffold of the pre-mRNA and could not be corrected by SRSF1. Thus, we examined if another ESE-dependent RBP could restore U1 snRNP recruitment to *AdML* Δ3′SS. In order to identify the ESE, we mutated three nucleotides in a single-stranded region in the 5′ exon of *AdML* Δ3′SS (Supplementary Figure S5E, the mutant is termed ΔESE), which did not significantly alter splicing of *AdML* ΔESE with intact 3′SS but drastically reduced that of *AdML* Δ3′SS ΔESE (Figure 5D – lanes 3 & 4). Blocking of U1 snRNP with U1 antisense morpholino oligonucleotide suppressed splicing of both *AdML* ΔESE and *AdML* ΔESE Δ3′SS (Supplementary Figure S5D – compare lanes 3, 4 with 8, 9).

ESE-finder (57) indicates that the region mutated in *AdML* Δ3′SS ΔESE (UCACUCU) is an *in vitro*-selected binding site for SR protein SRSF5 (alias SRp40). The two-RRM SR proteins (SRSF1 and SRSF5) exhibit similar RBD-dependent RNA binding and RNA structural remodeling mechanisms (49). Additionally, like SRSF1, SRSF5 is also activated upon phosphorylation of its RS domain (58). Accordingly, we mixed *AdML* Δ3′SS with the RNA binding domain (RBD) of SRSF5 (Supplementary Figure S5A) and U1 snRNP, and purified the complex by ion-exchange chromatography (Figure 5E). SDS PAGE indicated that high levels of U1 snRNP co-eluted with *AdML* Δ3′SS that is also bound by SRSF5-RBD (Figure 5F). We also mixed the same components with *AdML* Δ3′SS ΔESE and purified the complexes (Figure 5E). SDS PAGE of the fractions revealed lower levels of U1 snRNP co-eluting with the pre-mRNA (compare Figures 5F and 5G). A slight reduction in MBP-MS2 levels was also observed with *AdML* Δ3′SS ΔESE compared to *AdML* Δ3′SS; this is due to precipitation of a portion of the pre-mRNA complexes on the column, which was observed with the recruitment-defective pre-mRNAs throughout this study. For example, the level of eluted MBP-MS2 with WT *AdML*-MS2 in the presence of U1 snRNP (Figure 1B) was slightly greater than that with MS2-tagged *AdML* mutants (Figures 1C, D, E, F). To test if SRSF5-RBD stabilizes U1 snRNP primarily through direct interaction, we examined the interaction between GST-SRSF5-RBD and U1 snRNP by GST pull down assay (Supplementary Figure S5F). A weak interaction between U1 snRNP and GST-SRSF1-RBD was observed, but none was detected between GST-SRSF5-RBD and U1 snRNP. We also used another variant of U1 snRNP assembled with a truncated Sm B (1-174 a.a.) in the GST pull-down assay (Figure 5H). This variant exhibited a stronger interaction between SRSF1-RBD and U1 snRNP but none between SRSF5-RBD and U1 snRNP.

Overall, these results suggest that SRSF5 stabilizes U1 snRNP on *AdML* Δ3′SS without a detectable interaction with U1 snRNP likely through pre-mRNA structural remodeling.

### Structural remodeling of *AdML* variants by SR proteins correlates with differences in the global structural scaffold of the *AdML* variants

To correlate recruitment of early spliceosomal components and remodeling of *AdML*, we examined the SHAPE reactivity of *AdML* and its variants both in the protein-free state and in a complex with the RNA binding domain of two SR proteins, SRSF1 and SRSF5. A comparison of the SHAPE reactivity differential (Ad+SR1)-Ad (calculated by subtracting SHAPE reactivity of each nucleotide of mock-treated *AdML* from that of the corresponding nucleotide of SRSF1-RBD-bound *AdML*) and (Ad+U1)-Ad (calculated by subtracting SHAPE reactivity of each nucleotide of mock-treated *AdML* from that of the corresponding nucleotide of the U1 snRNP-bound *AdML*) is shown in Figure 6A. We anticipated that SRSF1-RBD might remove base-pairing constraints (49) from certain nucleotides that interact with U1 snRNP since the ternary SRSF1-RBD:*AdML*:U1 snRNP complex is more stable than the binary *AdML*:U1 snRNP complex. Henceforth, we looked for nucleotides that were more flexible in the SRSF1-RBD-bound state but less flexible in the U1 snRNP-bound state compared to protein-free *AdML*. This trend was distinctly observed in the 26^th^ and 43^rd^ nucleotides in the 5′ exon and six nucleotides in the intron (83^rd^, 87^th^, 102^nd^, 123^rd^, 124^th^, and 139^th^ nt) (marked with arrows in Figure 6A). The remainder of the nucleotides showing distinct changes exhibited a complex pattern. For example, there were nucleotides less flexible in both SRSF1-RBD-bound and U1 snRNP-bound RNAs, or less flexible in SRSF1-RBD-bound RNA but more flexible in U1 snRNP-bound RNA compared to those of the protein-free pre-mRNA. Additional work is needed to elucidate the mechanisms and functional consequences of these distinct changes.

**Figure 6.**
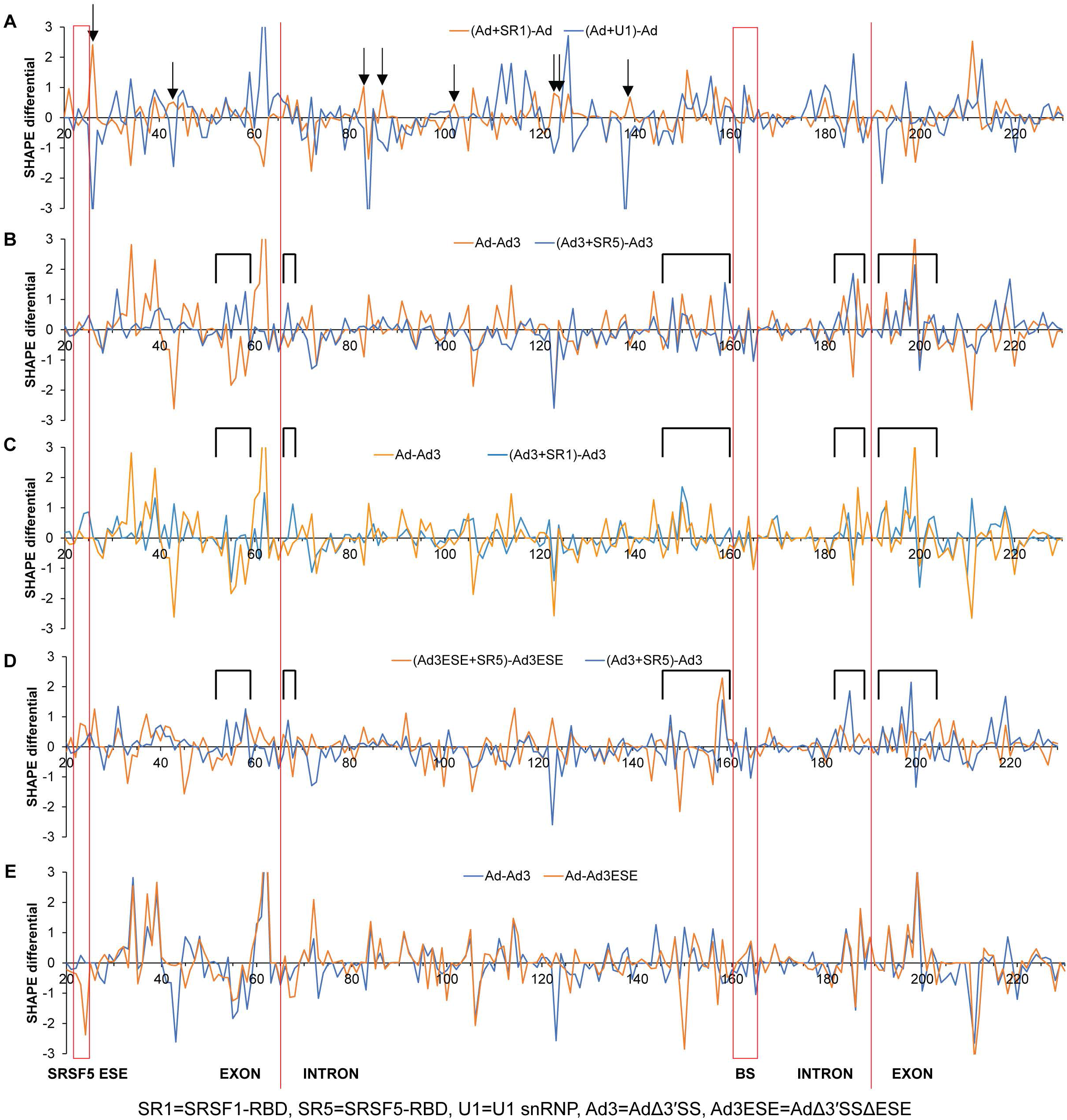
Structural remodeling of *AdML* variants by SR proteins correlates to difference in global structural scaffold of the pre-mRNA. (A) SHAPE differentials (Ad+SR1)-Ad (calculated by subtracting the SHAPE reactivity of each nucleotide of mock-treated *AdML* from that of *AdML*+SRSF1-RBD complex) and (Ad+U1)-Ad (differential of *AdML*+U1 snRNP complex and mock-treated *AdML*) are overlaid onto each other; nucleotides exhibiting a higher reactivity in *AdML*+SRSF1-RBD complex but a lower one in *AdML*+U1 snRNP complex compared to the protein-free *AdML* are marked with arrows; the positions of the SRSF5 ESE, the 5′ and the 3′ exon-intron junctions, and the BS are indicated with vertical lines. (B) SHAPE differentials Ad-Ad3 (differential of mock-treated *AdML* WT and *AdML* Δ3′SS) and (Ad3+SR5)-Ad3 (differential of *AdML* Δ3′SS+SRSF5-RBD complex and mock-treated *AdML* Δ3′SS) are overlaid onto each other; segments flanking the splice sites and upstream of the BS whose SHAPE reactivities are enhanced upon engagement of SRSF5-RBD to *AdML* Δ3′SS are marked with open rectangles. (C) SHAPE differentials Ad-Ad3 and (Ad3+SR1)-Ad3 (differential of *AdML* Δ3′SS+SRSF1-RBD complex and *AdML* Δ3′SS) are overlaid onto each other. (D) SHAPE differentials (Ad3ESE+SR5)-Ad3ESE (differential of *AdML* Δ3′SS ΔESE + SRSF5-RBD complex and mock-treated *AdML* Δ3′SS ΔESE) and (Ad3+SR5)-Ad3 are overlaid onto each other. (E) SHAPE differentials Ad-Ad3 and Ad-Ad3ESE are overlaid onto each other showing largely similar profiles.

As shown in Figures 6B-E, we compared SHAPE reactivity differentials of *AdML* variants and their complexes with the SRSF1-RBD (denoted by SR1 in shorthand) and SRSF5 (denoted by SR5). The comparison of the SHAPE reactivity differential of *AdML* and *AdML* Δ3′SS (Ad-Ad3) and (Ad3+SR5)-Ad3 revealed that nucleotides upstream of the BS and flanking 3’SS and the 5’SS become more reactive upon SRSR5-RBD binding to *AdML* Δ3′SS (Figure 6B, marked with open rectangles above the plot). An increase in reactivity/flexibility in these critical segments near the splice signals of *AdML* Δ3′SS upon binding of SRSF5-RBD could explain restoration of U1 snRNP binding to *AdML* Δ3′SS. Interestingly, some of these segments have lower SHAPE reactivity in *AdML* Δ3′SS compared to *AdML* WT. This suggests that binding of SRSF5-RBD to *AdML* Δ3′SS causes a reversal to WT SHAPE reactivity in these segments. The overall pattern of structural remodeling of *AdML* Δ3′SS by SRSF1-RBD is different from that by SRSF5-RBD. In particular, segments surrounding the 5′SS, BS and 3′SS are less reactive upon SRSF1-RBD binding (Figure 6C). The lack of reactivity in these critical regions of the substrate may explain why SRSF1 fails to facilitate recruitment of U1 snRNP (and likely other early spliceosomal components) to the *AdML* Δ3′SS substrate.

Finally, we examined the effect of SRSF5-RBD binding to *AdML* Δ3′SS ΔESE (Figure 6D). The increase in reactivity of the critical segments indicated above observed in *AdML* Δ3′SS upon binding of SRSF5-RBD was lost with the exception of the BS region. Overall, these results may explain why the splicing defect of *AdML* Δ3′SS ΔESE cannot be rescued by SRSF5-RBD despite SHAPE differentials Ad-Ad3 and Ad-Ad3ESE being largely comparable (Figure 6E).

We further carried out transfection-based *in vivo* pre-mRNA structure-probing to examine the *in vivo* implications of these findings. Cells with U1 snRNP blocked out with U1 snRNA antisense morpholino olinucleotide (U1 AMO) were transfected with *AdML* WT, *AdML* Δ3′SS, and *AdML* Δ3′SS ΔESE, and the SHAPE reaction was carried out with NAI (Methods). The splicing-competent substrates *AdML* WT and *AdML* Δ SS exhibited a greater level of flexibility across the pre-mRNA compared to *AdML* Δ3′SS ΔESE (Supplementary Figure S6A). This is in agreement with our previous observation that splicing competent pre-mRNAs are more unstructured *in vivo* (18). Additionally, the correlation between the moving averages (period = 6) of *in vivo* SHAPE reactivity of *AdML* WT and *AdML* Δ3′SS was significantly greater than those between *AdML* Δ3′SS & *AdML* Δ3′SS ΔESE and *AdML* WT & *AdML* Δ3′SS ΔESE (Supplementary Figure S6B). The structural difference between the *AdML* Δ3′SS variants *in vivo* is the functional consequence of the mutation in the SRSF5 ESE in the 5′ exon. This is because *AdML* Δ3′SS and *AdML* Δ3′SS ΔESE in the protein-free state did not exhibit a large structural variation (Figure 6E). Hence, this result highlights the role of SRSF5-mediated structural remodeling of *AdML* Δ3′SS for its splicing *in vivo*.

Overall, these results suggest that both the pattern of ESE-dependent pre-mRNA structural remodeling and its consequence for early spliceosome assembly efficiency are regulated by the global structural scaffold of *AdML* pre-mRNA.

### Interplay between ESE and the global structure of *AdML* variants regulates recruitment of U2AF65/U2AF35

We next examined whether binding of the 3′ end-recognizing factors U2AF65 and U2AF35 was affected by the above-described splice signal mutations. We mixed U1 snRNP, SF1_320_, U2AF65, and U2AF35 with *AdML* WT (bound to MBP-MS2), and performed anion-exchange chromatography (red line in Supplementary Figure S7A). From SDS PAGE analysis of the fractions, we did not observe clear binding of any splicing factor other than U1 snRNP (Supplementary Figure S7B), perhaps due to requirements of additional ESE-dependent splicing factors such as SR proteins. Accordingly, we added SRSF1 to the reaction mixture and purified the complexes by chromatography (blue line in Supplementary Figure S7A). SDS PAGE analysis of the fractions revealed a high level of U1 snRNP co-eluting with the MBP-MS2-bound pre-mRNA (Supplementary Figure S7C; raw SDS gel images are provided in Supplementary File 3); closer examination of the gel image revealed small (sub-stoichiometric) levels of U2AF65 and U2AF35 but not SF1_320_. In Supplementary Figure S7C and the subsequent SRSF1-containing gels, SRSF1 band intensity appeared higher than the other protein bands, since SRSF1 stains about five-times more intensely than U1-A as well as U1-70k (Supplementary Figure S7K).

Two evolutionarily conserved serine residues (S80/S82) of SF1 are heavily phosphorylated *in vivo*, and this phosphorylation is reported to enhance its interactions with the pre-mRNA and U2AF65 (59). Therefore, we used the phosphomimetic variant of SF1_320_ – SF1_320_ (S80E/S82E) in the chromatographic purification assay (Figure 7A – blue line) to test if this enhanced binding of SF1_320_, U2AF65, and U2AF35, but it was undetectable (Figures 7B and Supplementary Figure S7C). Next, we examined formation of this complex with the *AdML* ΔPPT (Figure 7A – grey line) followed by SDS PAGE analysis (Figure 7C) to test if this complex is specific. We reproducibly observed an absence of the U2AF65 and U2AF35 protein bands. Although SRSF1 enhanced the levels of bound U1 snRNP to *AdML* ΔPPT (Figure 1F), it was not as robust as with *AdML* WT; the peak of U1 snRNP proteins co-eluted with *AdML* WT complexes (a gradual increase between Fr 25 – 27 and decrease between Fr 28-33 in Figure 7B) was not observed with *AdML* ΔPPT. We then tested the assembly of this complex with *AdML* Δ5′SS (Figures 7A – red line), which behaved like *AdML* ΔPPT (Figure 7D). Recruitment of U2AF65 and U2AF35 to the *AdML* Δ5′SS-GU mutant also appeared to be strongly diminished (Supplementary Figure S7D, S7E). We also tested the recruitment of U2AF65 and U2AF35 to *AdML* Δ3′SS in the presence of SRSF5-RBD by chromatography (Figure 7E) followed by SDS PAGE of the fractions (Figure 7F), which revealed a WT-level recruitment of U2AF65 and U2AF35. We have not examined the identity of the SRSF5-enhancer, likely near the 3′SS, which promotes recruitment of U2AF65 and U2AF35 to *AdML* Δ3′′

**Figure 7.**
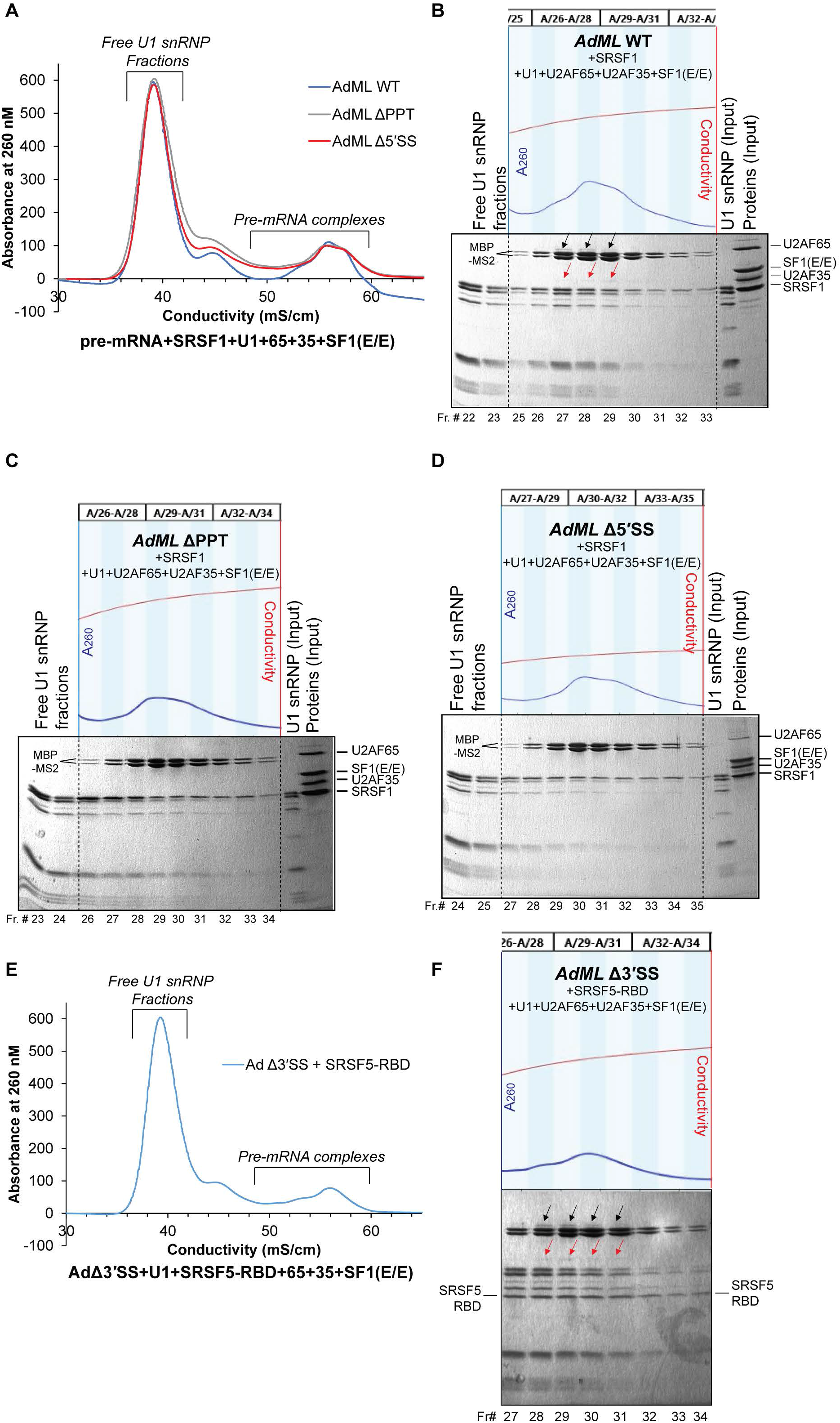
Interplay between ESE and global structure of *AdML* variants regulates recruitment of U2AF65/U2AF35. (A) Chromatograms of purification of *AdML* complexes formed with WT, Δ5′SS, and ΔPPT in the presence of U1 snRNP, U2AF65, U2AF35, SF1_320_ (S80E/S82E), and the ESE-dependent RBP SRSF1; the raw chromatograms including fraction numbers are shown above each gel. (B, C, D) SDS PAGE of fractions corresponding to chromatographically purified *AdML* WT complexes (B), ΔPPT complexes (C), Δ5′SS complexes (D); ‘black arrow’ indicates protein bands of U2AF65 and ‘red arrow’ U2AF35; the chromatography fractions marked as pre-mRNA complexes in A are formed with MBP-MS2-bound pre-mRNAs and concentrated by amylose pull-down before SDS PAGE; fraction numbers are given under each SDS gel; raw SDS gel images are provided in Supplementary File 3. (E) Chromatogram of purification of complexes formed with *AdML* Δ3′SS, U1 snRNP, SRSF5-RBD, U2AF65, U2AF35, and SF1_320_ (S80E/S82E). (F) SDS PAGE of fractions containing the pre-mRNA complexes corresponding to the chromatogram shown in E; the positions of U2AF65 and U2AF35 bands are marked with arrows as in B; the position of SRSF5 band is indicated.

Since we did not detect SF1_320_ in the purified *AdML* WT complexes, we asked if SF1_320_ is required for binding of U2AF65 and U2AF35 to the WT pre-mRNA. We carried out similar chromatographic experiments with *AdML* WT with all components except for SF1_320_ (Supplementary Figure S7F). Co-elution of U2AF65 or U2AF35 with the pre-mRNA (Supplementary Figure S7G) was less robust than in the presence of SF1_320_ (Supplementary Figure S7C). Additionally, neither chromatographic (Supplementary Figures S7H, S7I) nor amylose-pull-down assays (Supplementary Figure S7J) could detect SF1_320_ or SF1_320_ (S80E/S82E) bound to the pre-mRNA in the absence of U2AF65 and U2AF35. These results suggest that in our simplified system consisting of purified components, SF1_320_ is required for stabilization of U2AF65 and U2AF35 on *AdML* but is not retained in the complex suggesting it may have a chaperone-like functionality. A previous report indicated that SF1 may even be skipped for several mammalian splicing events in a splicing-competent nuclear extract containing all nuclear proteins (60).

Overall, mutations in non-cognate splice signals could impede ESE-dependent recruitment of U2AF65 and U2AF35 to *AdML* likely through disruption of the global RNA structural environment.

## DISCUSSION

The efficiency of a splicing event depends not only on the nucleotide sequence of the splicing motifs (i.e. splice signals and SREs) but also on their context (25). Due to this context-dependent functionality of splicing motifs, the regulatory potential of a splicing factor is not fully correlated to the level of conservation within its cognate binding motif. In the current work, we examined the potential of the global structural scaffold of a model pre-mRNA substrate – *AdML* – to regulate the engagement of early spliceosomal components and the functionalities of RBPs, which bind the splice signals and the SREs, respectively. We used SHAPE, FRET, chromatographic binding assays, and transfection-based splicing assays to examine the regulation of splice signal recognition by the global structural scaffold of the pre-mRNA. Our work suggests that the global structural scaffold of *AdML* provides the context to the splicing motifs by integrating/embedding them into a structured splicing unit. This conclusion is derived from mutational studies which show that splice signal mutations disrupt its global structure and diminish binding of U1 snRNP known to recognize the 5′SS. These results suggest that U1 snRNP recognizes the structural scaffold of *AdML* involving all splice signals. Similarly, U2AF65 and U2AF35 also failed to recognize *AdML* splice signal variants, again pointing to the importance of a folded structure in splicing factor recruitment. FRET experiments support the structural scaffold model, which reveal spatial proximity of segments far apart in the primary sequence. This structural framework could also stop pseudo-splicing motifs from being recognized. Our result obtained with a human adenovirus 2 pre-mRNA conforms to the earlier observation that 5 SS and the BS are spatially proximal in a protein-free yeast pre-mRNA (61).

In addition to the splice signals, the functionalities of SREs of *AdML* could also be regulated by its global structural scaffold. We observed that the *AdML* Δ3′SS, which splices efficiently *in vivo* from a cryptic 3′SS 6-nt downstream of the authentic 3′SS, cannot recruit U1 snRNP in the protein-free state unlike *AdML* WT but can do so in the presence of the RNA binding domain of SRSF5. SHAPE probing suggests that SRSF5-RBD binding can induce reversal of certain *AdML* Δ3′SS-specific structural features to *AdML* WT-like features. We further observed that binding of SRSF1-RBD to *AdML* Δ3′SS does not induce such reversal of SHAPE reactivities as efficiently as SRSF5-RBD and does not enable recruitment of U1 snRNP. In contrast, SRSF1-RBD stabilizes the U1 snRNP complex formed with WT *AdML*. These results led us to propose two modes of cooperation between the pre-mRNA structural scaffold and the ESE (Figure 8). The pre-mRNA with a favorable global structure could engage U1 snRNP (or other early spliceosomal components), with further stabilization by ESE-dependent RBPs for assembly of the early spliceosome. The structure of some pre-mRNAs may not be favorable to recruit U1 snRNP or other early spliceosomal components and could do so only upon structural remodeling by RBPs in an ESE-dependent manner. The earlier observations that SREs could enable alteration of the interactions within U1 snRNP-dependent pre-mRNA complexes (62,63) also potentially highlight the interplay between the global structural scaffold and SRE-dependent pre-mRNA structural remodeling in the regulation of early spliceosome assembly. Nonetheless, it will be intriguing to investigate the prevalence of regulation of functionalities of splice signals and SREs by the global three-dimensional structure of the pre-mRNAs across the transcriptome of different cell types. This insight might prove to be highly useful for explaining the regulation of alternative splicing, in characterizing the effect of disease-causing mutations, and in identifying methods to alter splicing efficiency for therapeutic purposes.

**Figure 8.**
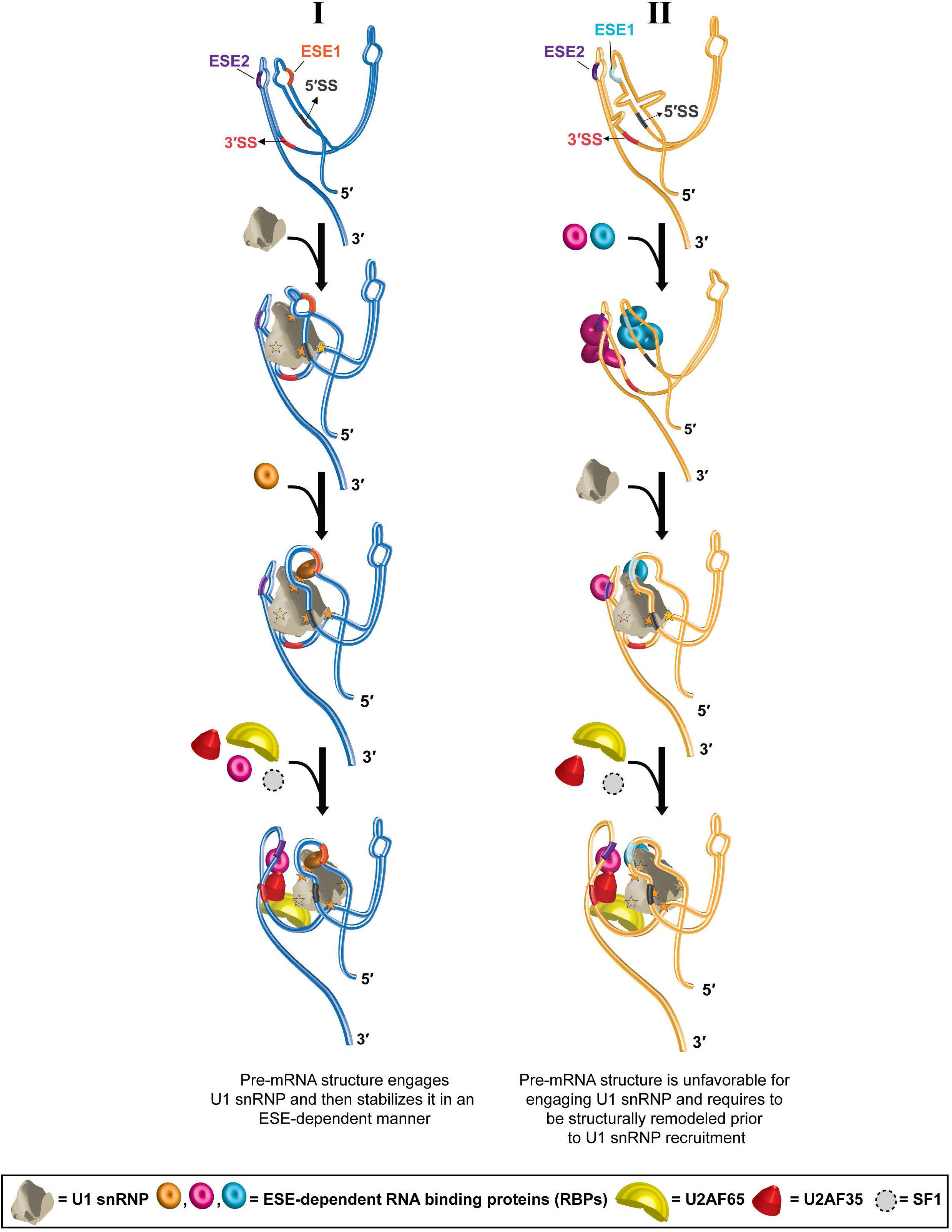
Proposed model of splicing substrate activation by cooperation between global pre-mRNA structure and ESEs. **Pathway I:** The global three-dimensional structure of a pre-mRNA enables engagement (strong or weak) of U1 snRNP; the splice sites and ESEs in the 5′ and 3′ exons are indicated; PPT and BS immediately upstream of the 3′ are not drawn for clarity. Four hypothetical contact points between U1 snRNP and the protein-free pre-mRNA including the one at 5′SS region are shown with yellow stars; the solid stars suggest contact on the top surface and the translucent stars the opposite surface. Recruitment of an RBP at an ESE of the 5′ exon further stabilizes U1 snRNP on the pre-mRNA – the additional contact point between the RBP and U1 snRNP is shown with a star. This complex has the right conformation to receive U2AF65 and U2AF35, which are further stabilized by an RBP bound at an ESE in the 3′ exon. SF1 may play a chaperone-like role in the latter process. **Pathway II:** The pre-mRNA structure is unfavorable for engagement of U1 snRNP and needs to be structurally remodeled by additional factors such as ESE-dependent RBPs. The ESE-dependent RBPs could bind the pre-mRNA in multiple copies cooperatively, which structurally remodels the pre-mRNA (18), enabling recruitment of U1 snRNP (the contact points between U1 snRNP and the pre-mRNA and that between U1 snRNP and the ESE-bound RBP are indicated with five stars). This complex then may receive U2AF65 and U2AF35.

The splice site recognition mechanism *in vivo* across the transcriptome is complex. The order of splicing of introns of highly varied lengths may not follow the order of intron synthesis (64) involving co-transcriptional or post-transcriptional splicing (65). Our model presumes that both 5′- and 3′-SS are present in the pre-mRNA during substrate definition, which could be true for post-transcriptional splicing but not co-transcriptional splicing. Due to the complexity of the spliceosomal system *in vivo*, it may be difficult to put forth a universal mechanism for recognition of splice sites across the transcriptome. However, our data and its extrapolations support a model that correlates with several previously described molecular recognition events related to co-transcriptional splicing. Context-dependent binding of splicing factors to full-length pre-mRNAs observed *in vitro* correlates with their *in vivo* binding (66). Additionally, many splicing factors, including U1 snRNP, are found to be associated with RNA polymerase II (RNAPII) (67,68). The U1 snRNP particle seems to be associated with both the 5′SS and the elongating RNAPII molecule at least until the following exon synthesis is completed (67). This might provide U1 snRNP the opportunity to recognize the splicing substrate involving both splice sites across an intron (or an exon). Looping out of the longer introns, which brings the splice sites closer, is also reported to enhance splicing (69) possibly by allowing close-knit association of splice signals and flanking segments within a three-dimensional structural framework of pre-mRNA for functional recruitment of the U1 snRNP. The physical separation of exons and introns between nuclear speckles and the nucleoplasm, and the presence of U snRNPs at the periphery of nuclear speckles (70) could also support cross-intron and/or cross-exon recognition of the pre-mRNA by U1 snRNP.

There are many additional questions that need to be answered to fully comprehend the impact of the global structural scaffold of the pre-mRNA on splicing. Both exon- and intron-definition of a pre-mRNA is important to explain the fidelity of splicing (67), and the early spliceosome could assemble across the intron as well as across the exon (26). Additionally, earlier observations suggest that the polypyrimidine-tract binding protein (PTB) prevents intron-definition but does not block U1 snRNP recruitment at the upstream 5′SS (71). Thus, it would be interesting to examine if the structural framework recognized by U1 snRNP may be formed across the internal exons and how PTB and other splicing suppressors interact with the pre-mRNA global structural landscape. Furthermore, a previous study of a substrate with two alternative 5′SSs suggested that both 5′SSs are engaged by U1 snRNP in the early spliceosome (E-complex) and one is purged during ATP-dependent transition to the A-complex (72). Determining whether the pre-mRNA global structure could mediate such alternative splice site selection at this early stage requires further investigation. The distance of an alternative splice site from the exon-intron junctions often impacts its selection efficiency. However, the mechanism for how the proximal site is favored in some cases while the distal one is favored in others is currently unclear (73,74). Future investigations are needed to elucidate if this effect is regulated by the global structural scaffold of the pre-mRNA and its modulation. Moreover, transcriptome-wide binding studies suggest that human U2AF65 avoids the majority of non-authentic PPT-like sequences (75). Understanding how this is achieved by a coordinated action of DEK (76), hnRNP A1 (77), the global RNA structural scaffold, and other PPT-like sequence binding proteins that might occlude the unauthentic sites will also require further investigation. Additionally, recruitment of U2AF65 at alternative PPTs within the same intron (75), at the authentic PPT in coordination with other PPT-binding protein factors (78), or at the authentic PPT with the help of multiple exonic and intronic splicing enhancers (79) have also been reported. If and how these phenomena are correlated to the pre-mRNA structural scaffold requires further investigation. Moreover, the ability of U1 snRNP to recognize a structural landscape beyond the 5′SS-like sequence could be explored for understanding its binding mechanism to non-coding RNAs (80).

Overall, the results described above suggest that splicing motifs can be structurally and functionally integrated within a splicing-conducive three-dimensional pre-mRNA structural scaffold. Therefore, it may not be possible to comprehensively assign splicing code to individual sequences without considering their presentation within the pre-mRNA scaffold. This work provides the initial basis for understanding the contribution of the pre-mRNA global structural framework to the mammalian splicing code.

## DATA AVAILIBILITY

### Supplementary File 1

SHAPE reactivity of *AdML* variants obtained *in vitro* with 2 and 8 mM NMIA and *in vivo* with 100 mM NAI.

### Supplementary File 2

Rep 1, 2, 3: Three replicates of fluorescence emission obtained upon excitation with 550 nm light with *AdML* hybridized to DNA #2 and DNA #3, DNA #5, or DNA #6.

Disrupt: Fluorescence emission obtained with *AdML* hybridized to the structure disrupting 40-nt long DNA with the same combination of fluorescently labeled DNA probes.

Low Mg: Fluorescence emission obtained with 0.1 mM MgCl_2_ instead of the usual 2 mM.

### Supplementary File 3

Raw gel images (.scn files) obtained with Bio-Rad Chemidoc pertaining to Figures 5F, 5G, 7B, 7C, 7D, 7F, Supplementary Figures S7B, S7C, S7E, S7G, S7I to be analyzed with the Image Lab software (Bio-Rad).

### Accession number

Raw and processed files pertaining to the RNA-seq data have been deposited to Gene Expression Omnibus under accession number GSE173178.

## AUTHOR CONTRIBUTION

KS designed and performed majority of the experiments, analyzed the data, and wrote the paper. TB designed and performed EM experiments and wrote the paper. MF performed some protein purification experiments and critiqued the paper. SJ supervised FRET experiments. GG supervised the project and wrote the paper.

