## Supplementary Figures for "Discovery of a pre-mRNA structural scaffold as a contributor to the mammalian splicing code"

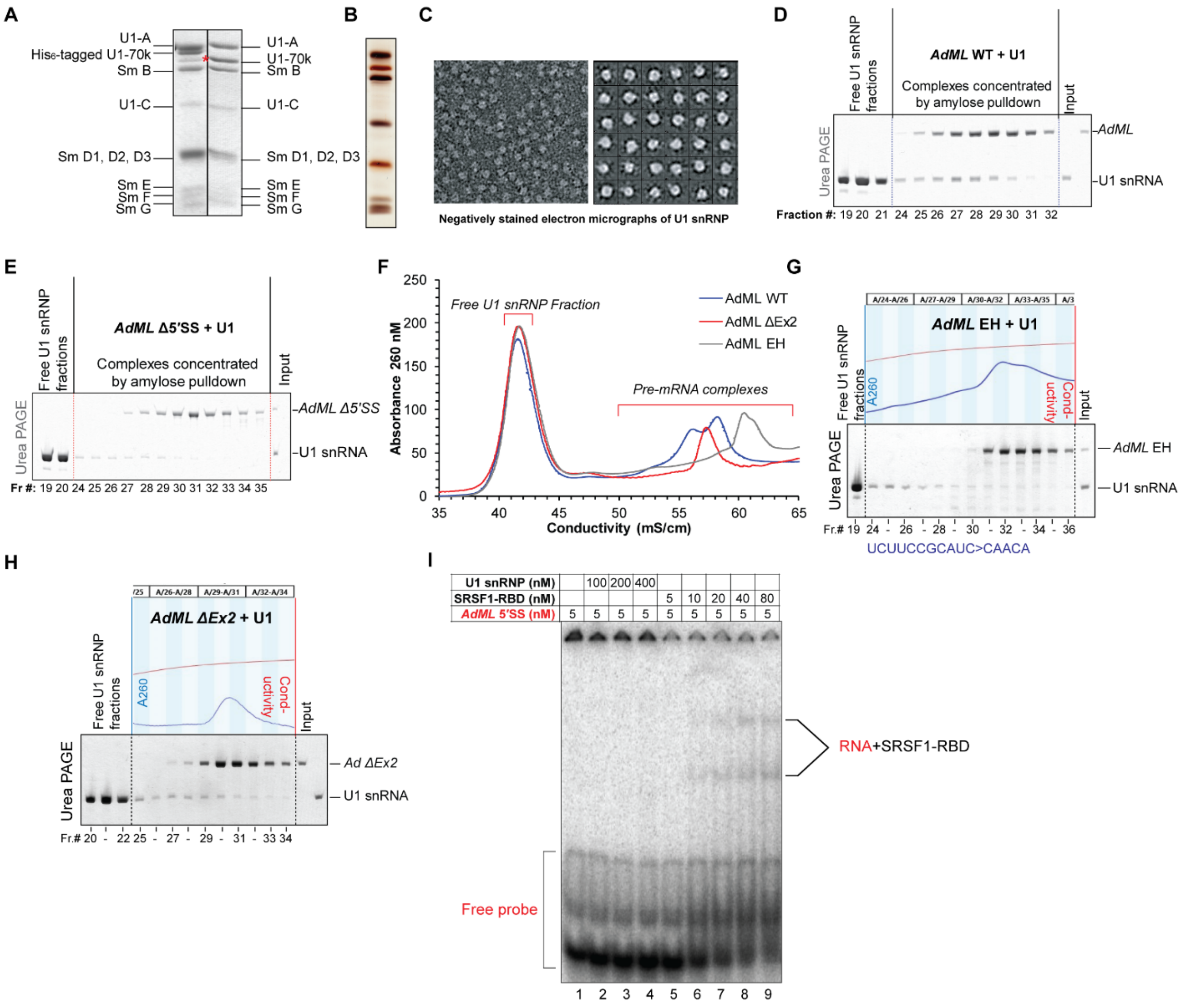


**Supplementary Figure S1. U1 snRNP binding to protein-free RNA**

(A) SDS PAGE of purified U1 snRNP assembled with His$_6$-tagged U1-70k (left) and U1-70k with cleaved His$_6$-tag (right); '*' indicates naturally proteolyzed U1-70k that co-migrates with His$_6$-cleaved U1-70k; U1-C stains weakly with Coomassie blue. (B) Silver-stained purified U1 snRNP showing appropriate staining of U1-C and weaker staining of U1-70k. (C) Negatively stained electron micrograph of purified U1 snRNP exhibiting high homogeneity of particles (left); a preliminary 2D-average analysis of 2527 particles into 48 classes (only 36 shown) exhibiting the distinctive Sm core and additional protuberances (right). (D & E) Urea gel of fractions of WT *AdML* (D) and *AdML* Δ5′SS (E) complexes corresponding to the chromatogram shown in Figure 1A; the fractions indicated as 'pre-mRNA complexes' in the chromatogram are concentrated by amylose pull-down; lanes labeled as 'free U1 snRNP fractions' contain RNA extracted from 0.1 ml of each of the 0.3 ml of fractions within the peak marked with the same name in the chromatogram shown in Figure 1A; 2 pmol U1 snRNA & 0.5 pmol pre-mRNA are used as input in the urea-gel; corresponding fraction numbers are given below each lane. (F) Chromatograms of purification U1 snRNP-dependent complexes formed with *AdML* EH and *AdML* ΔEx2 mutants; chromatogram of purification of WT *AdML* complexes are shown for comparison; *AdML* EH mutant lacks 3X MS2 binding sequence at its 3′ end. (G & H) Urea PAGE analysis of the total RNA contents corresponding to purification chromatograms shown in F – *AdML* EH (G) and *AdML* ΔEx2 (H); the raw chromatogram representing the fractions containing the pre-mRNA complexes are shown above each gel, where blue line (absorbance trace) and red line (conductivity trace) are placed along primary and secondary *y*-axes, respectively. (I) EMSA showing a lack of binding of U1 snRNP (lanes 2, 3, 4) and binding of SRSF1-RBD (lanes 5-9) to radiolabeled 14-nt long *AdML* 5′SS RNA; the red script indicates radiolabeled components.

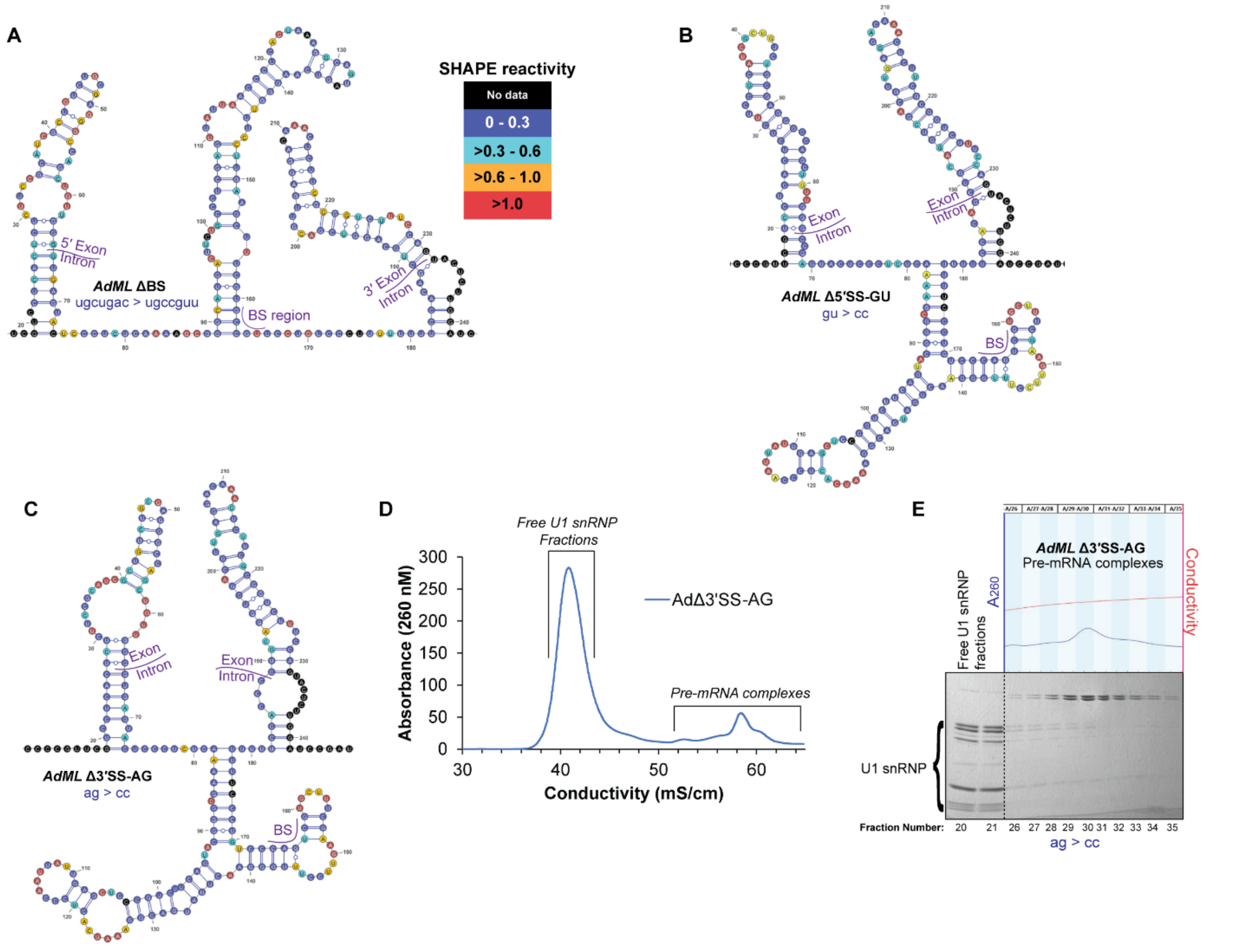


F

| AdΔ3_K1 | AdΔ3_K2 | AdΔ5_K1 | AdΔ5_K2 | AdΔBS_K1 | AdΔBS_K2 | AdWT_K1 | AdWT_K2 |
|---|---|---|---|---|---|---|---|
| 3907.56 | 4701.2 | 4093.36 | 4879.07 | 9569.58 | 10403.7 | 5865.63 | 6703.63 |
| | 4698.53 | | 4904.51 | | 10419.4 | | 6719.82 |
| | 4686.43 | | 4879.17 | | 10427.3 | | 6705.12 |
| | 4717.14 | | 4903.46 | | 10416.1 | | 6695.83 |
| | 4703.7 | | 4879.16 | | 10398.7 | | 6725.42 |
| | 4691.14 | | 4911.67 | | 10423.4 | | 6698.43 |
| | 4692.47 | | 4881.89 | | 10416 | | 6722.99 |
| | 4702.06 | | 4882.45 | | 10409.8 | | 6723.42 |
| | 4701.18 | | 4874.34 | | 10417.7 | | 6711.15 |
| | 4692.46 | | 4874.93 | | 10410 | | 6696.28 |

**Supplementary Figure S2. SHAPE-derived secondary structure models of *AdML* mutants** (A, B, C) SHAPE-derived secondary structure models of *AdML* ΔBS, *AdML* Δ5′SS-GU, and *AdML* Δ3′SS-AG; the nucleotides are color-coded according to their SHAPE reactivity as indicated in the associated legend; the mutated sequences are shown. (D, E) Assessment of U1 snRNP binding to *AdML* Δ3′SS-AG mutant by chromatographic purification of pre-mRNA+U1 snRNP reaction mixture (D) and subsequent SDS PAGE of the fractions showing highly weakened U1 snRNP binding (E). (F) DREEM-derived Bayesian Information Criteria (BIC) score of the K1 class and ten K2 classes showing a lower BIC score for the K1 class for each of the *AdML* variants.

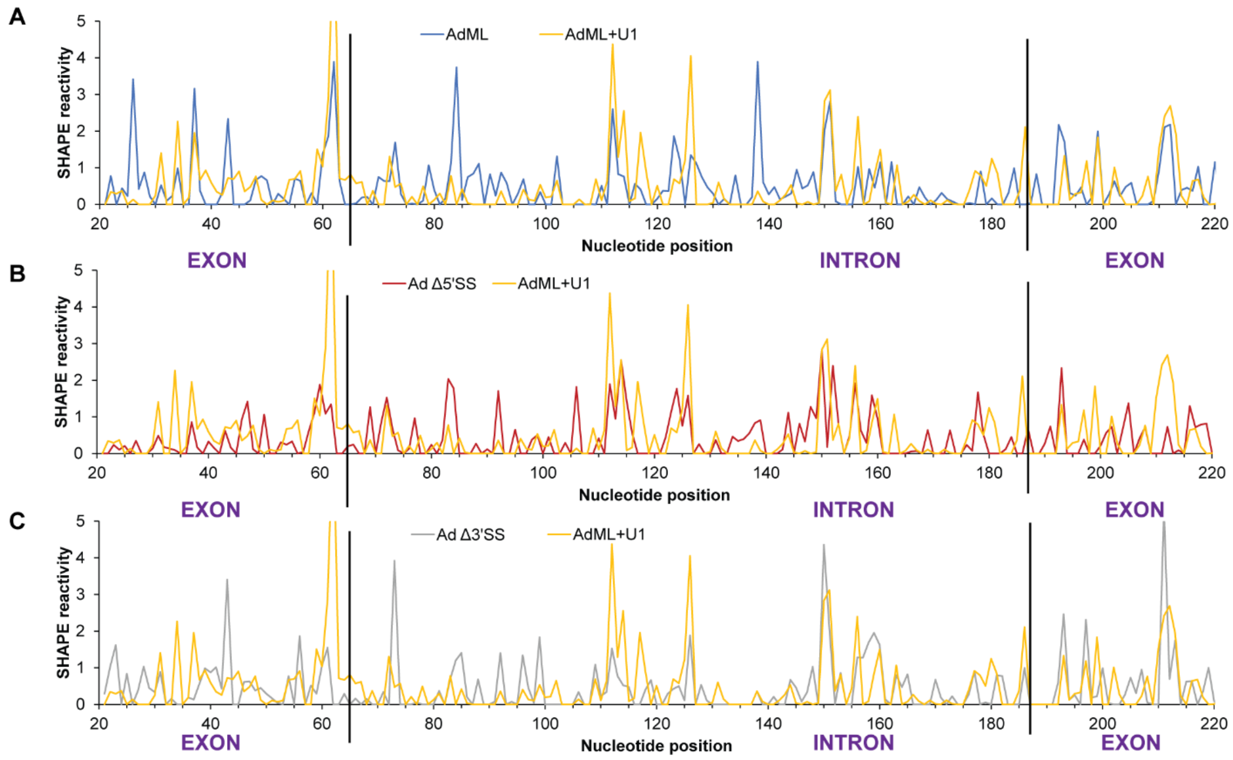


**Supplementary Figure S3. Comparison of SHAPE reactivity of *AdML* WT, Δ5′SS, Δ3′SS with that of *AdML* + U1 snRNP complex**

(A, B, C) SHAPE of reactivity of *AdML* WT (A), *AdML* Δ5′SS (B), and *AdML* Δ3′SS (C) obtained with 2 mM NMIA are overlaid onto that of *AdML* + U1 snRNP complex; the exon-intron boundaries are indicated with black vertical lines.

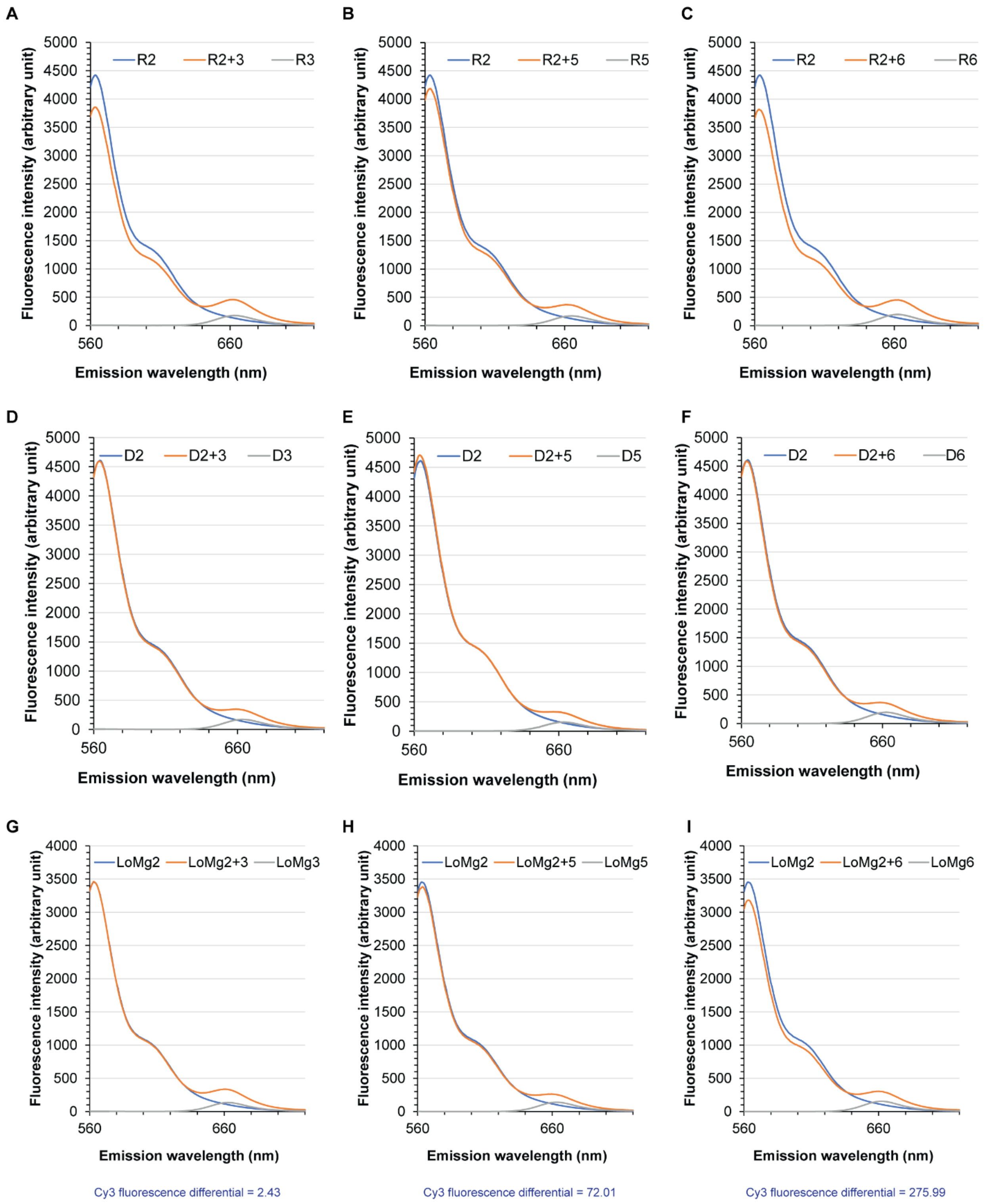


**Supplementary Figure S4. Spatial proximity of *AdML* helices tested by FRET**

(A) Representative plot demonstrating fluorescence-emission upon excitation at 550 nm from Cy3- and Cy5-DNA probes annealed to folded WT *AdML*: with annealed Cy3-probe (DNA #2) alone (blue line), annealed Cy5-probe (DNA #3) alone (grey line), and annealed both probes (DNA #2 & #3) (orange line). (B & C) Representative plots of similar emission as shown in A from probe-pairs #2 and #5 (B) and #2 and #6 (C). (D, E, F). Fluorescence-emission from the same probe-pairs as in A, B, C annealed to *AdML* WT with its structure disrupted by hybridization to the 40-nt long DNA indicated with a red line in Figure 4A. (G, H, I) Fluorescence-emission from the same probe-pairs as in A, B, C annealed to *AdML* WT under low $Mg^{2+}$ condition (0.1 mM $MgCl_2$ instead the usual of 2 mM); the difference between Cy3 fluorescence values in the presence and the absence of the Cy5 primer is shown below each plot.

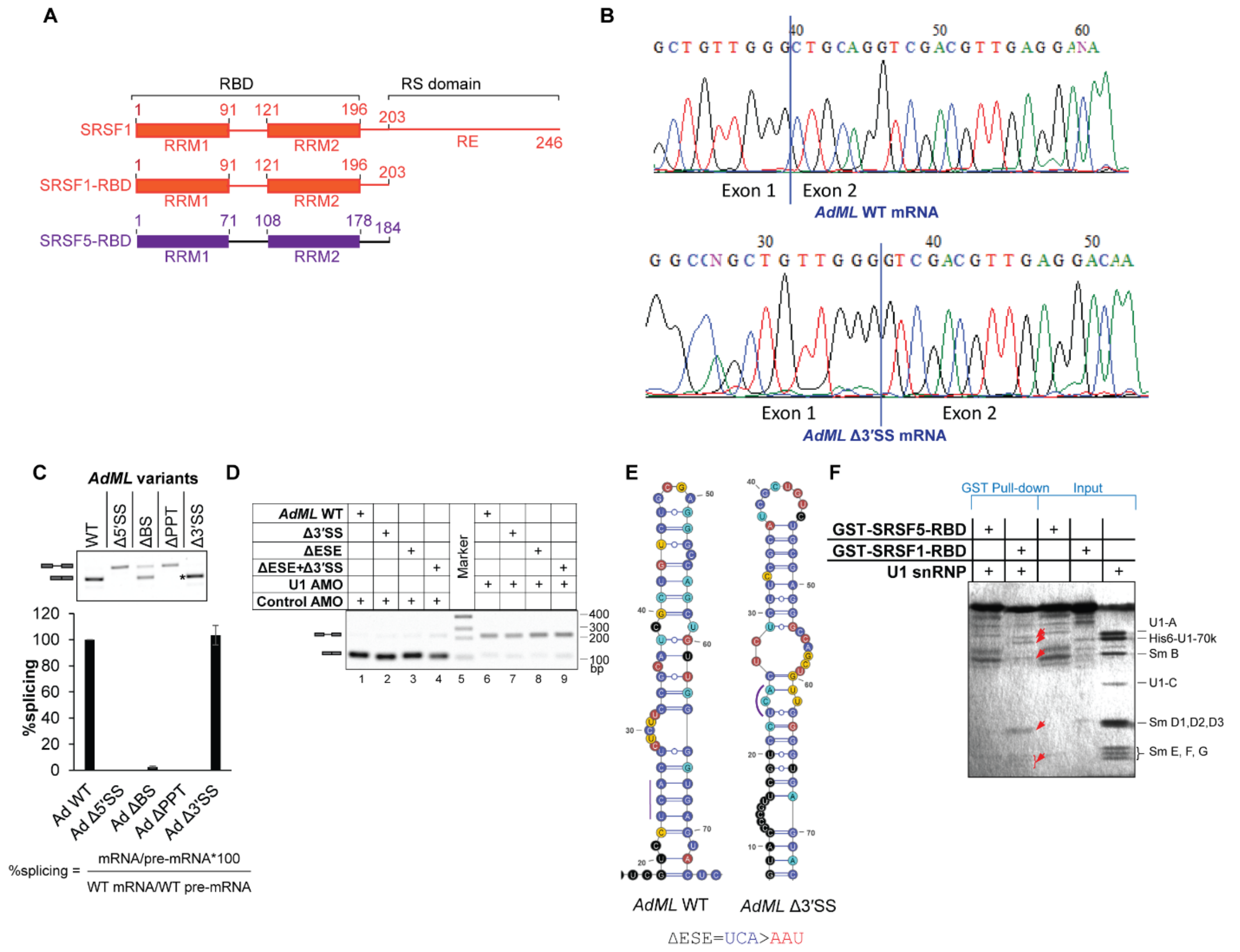


**Supplementary Figure S5. Mutation in splicing motifs and functionalities of SR proteins**
(A) SR protein constructs used in this study; the serine residues in the C-terminal RS domain of SRSF1 are replaced with glutamate (RE) generating the phosphomimetic variant (RBD = RNA binding domain); SRSF5 construct represents its RBD only. (B) Sequencing chromatogram of *AdML* mRNA produced by authentic splicing (top) and *AdML* Δ3′SS mRNA produced by cryptic splicing from a 3′SS 6-nt downstream of the authentic 3′SS (bottom); exon-exon junctions are indicated with blue vertical lines. (C) (top) Transfection-based splicing assay of WT *AdML* and its splice signal mutants; '*' indicates cryptic splicing product; (bottom) %splicing calculated from densitometric estimation of mRNA and pre-mRNA bands using the indicated formula of biological replicas; error bar indicates standard deviation (n = 3). (D) Transfection-based splicing assay of *AdML* variants in the presence of antisense morpholino oligonucleotide (AMO) complementary to the 5′ end of U1 snRNA or scrambled AMO. (E) SHAPE-derived secondary structure models of the 5′ exon of *AdML* WT and *AdML* Δ3′SS showing the location and sequence of mutation in *AdML* ΔESE. (F) GST-pull down assay showing the absence of detectable interaction between U1 snRNP and GST-SRSF5 (lane 1) and the presence of interaction between U1 snRNP and GST-SRSF1-RBD (lane 2); pulled down U1 snRNP proteins in lane 2 are marked with red arrows.

A

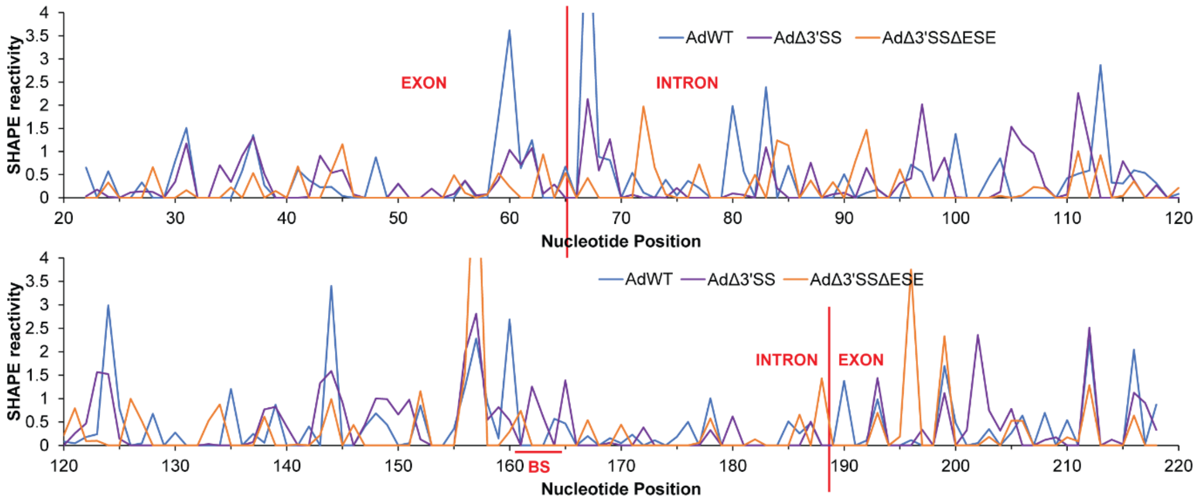


B

**Correlation Table**

| | |
|---|---|
| *AdML* WT vs *AdML* Δ3′SS | r = 0.55<br>*p* = 8.0E-17 |
| *AdML* WT vs *AdML* Δ3′SS ΔESE | r = 0.30<br>*p* = 2.9E-05 |
| *AdML* Δ3′SS vs *AdML* Δ3′SS ΔESE | r = 0.34<br>*p* = 1.1E-06 |

**Supplementary Figure S6. *In vivo* SHAPE reactivity of *AdML* variants in cells with U1 snRNA hybridized to U1 AMO**

(A) SHAPE reactivity of *AdML* WT (blue line), *AdML* Δ3′SS (violet line), and *AdML* Δ3′SS ΔESE (orange line); the plot is split into two segments – $20^{th}$ - $120^{th}$ nt (top) and $120^{th}$ - $220^{th}$ nt (bottom). (B) Correlation coefficient (r) and the corresponding *p* value for the moving averages (period = 6) of SHAPE reactivities of the *AdML* variants showing a greater correlation between the two splicing competent substrates *AdML* WT and *AdML* Δ3′SS.

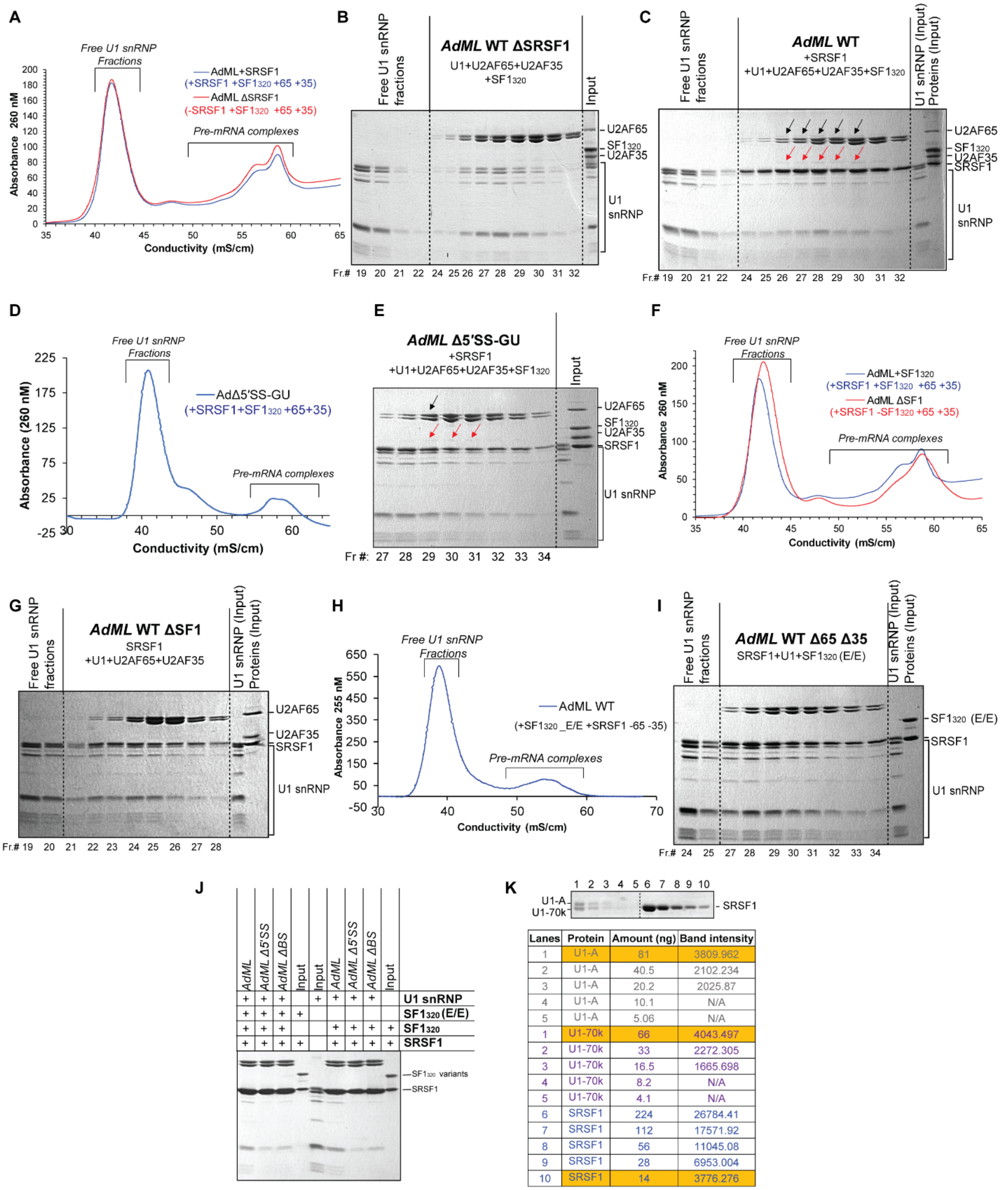


| Lanes | Protein | Amount (ng) | Band intensity |
|---|---|---|---|
| 1 | U1-A | 81 | 3809.962 |
| 2 | U1-A | 40.5 | 2102.234 |
| 3 | U1-A | 20.2 | 2025.87 |
| 4 | U1-A | 10.1 | N/A |
| 5 | U1-A | 5.06 | N/A |
| 1 | U1-70k | 66 | 4043.497 |
| 2 | U1-70k | 33 | 2272.305 |
| 3 | U1-70k | 16.5 | 1665.698 |
| 4 | U1-70k | 8.2 | N/A |
| 5 | U1-70k | 4.1 | N/A |
| 6 | SRSF1 | 224 | 26784.41 |
| 7 | SRSF1 | 112 | 17571.92 |
| 8 | SRSF1 | 56 | 11045.08 |
| 9 | SRSF1 | 28 | 6953.004 |
| 10 | SRSF1 | 14 | 3776.276 |

**Supplementary Figure S7: Analysis of recruitment of SF1, U2AF65, and U2AF35 to *AdML***
(A) Chromatograms of purification of complexes formed with WT *AdML*, U1 snRNP, U2AF65, $SF1_{320}$, and U2AF35 with (blue line) and without (red line) SRSF1. (B) SDS PAGE of chromatographic fractions corresponding to the red line in the chromatogram shown in (A); pre-mRNA complexes were concentrated by amylose pull-down; fraction numbers are indicated; raw SDS gel images of chromatographic fractions are provided in Supplementary File 3. (C) SDS

PAGE of chromatographic fractions corresponding to the blue line in the chromatogram shown in (A); 'black arrow' indicates bands of U2AF65 and 'red arrow' U2AF35. (D) Chromatogram of purification of complexes formed with *AdML* Δ5′SS-GU, U1 snRNP, SRSF1, U2AF65, U2AF35, and $SF1_{320}$. (E) SDS PAGE of fractions representing pre-mRNA complexes in D. (F) Chromatogram representing purification of complexes formed with WT *AdML*, U1 snRNP, SRSF1, U2AF65, and U2AF35 in the presence (blue line) or the absence of $SF1_{320}$ (red line). (G) SDS PAGE of chromatographic fractions corresponding to the red line in (F). (H) Chromatogram of purification of complexes formed with *AdML*, U1 snRNP, SRSF1, and $SF1_{320}$ (S80E/S82E) in the absence of U2AF65 and U2AF35. (I) SDS PAGE of fractions corresponding to the chromatogram in H. (J) Amylose pull-down assay (without chromatographic separation) showing absence of $SF1_{320}$ as well as $SF1_{320}$ (S80E/S82E) in the complex formed with *AdML*, *AdML* Δ5′SS, and *AdML* ΔBS in the presence of U1 snRNP and SRSF1. (K) Comparison of staining property of SRSF1 to those of U1-A and U1-70k by Coomassie blue; the mass of protein and the densitometric values of band intensities are given in the table; 0.5, 1, 2, 4, and 8 pmol of full-length phosphomimetic SRSF1 (27.2 kDa) and 0.1625, 0.325, 0.65, 1.3, and 2.6 pmol of U1 snRNP resolved on an SDS gel was stained with Coomassie blue;  of U1 snRNP components, U1-A (31.25 kDa) and U1-70k (25.5 kDa) are shown; bands representing 2.6 pmol of U1-A (i.e. 81 ng U1-A) as well as 2.6 pmol of U1-70k (i.e. 66 ng U1-70k) have similar staining intensity as 0.5 pmol SRSF1 (i.e. 14 ng SRSF1).
